# Divergent evolution of a protein-protein interaction revealed through ancestral sequence reconstruction and resurrection

**DOI:** 10.1101/2020.03.27.012369

**Authors:** Louise Laursen, Jelena Čalyševa, Toby J. Gibson, Per Jemth

## Abstract

The postsynaptic density extends across the postsynaptic dendritic spine with Discs large (DLG) as the most abundant scaffolding protein. DLG dynamically alters the structure of the postsynaptic density, thus controlling the function and distribution of specific receptors at the synapse. PDZ domains make up one of the most abundant protein interaction domain families in animals. One important interaction governing postsynaptic architecture is that between the PDZ3 domain from DLG and cysteine-rich interactor of PDZ3 (CRIPT). However, little is know regarding functional evolution of the PDZ3:CRIPT interaction. Here, we subjected PDZ3 and CRIPT to ancestral sequence reconstruction, resurrection and biophysical experiments. We show that the PDZ3:CRIPT interaction is an ancient interaction, which was present in the last common ancestor of Eukaryotes, and that high affinity is maintained in most extant animal phyla. However, affinity is low in nematodes and insects, raising questions about the physiological function of the interaction in species from these animal groups. Our findings demonstrate how an apparently established protein-protein interaction involved in cellular scaffolding in bilaterians can suddenly be subject to dynamic evolution including possible loss of function.

## Introduction

The postsynaptic density extends across the postsynaptic dendritic spine and is composed of receptors, signaling enzymes, cytoskeletal structural elements and cytoplasmic scaffolding proteins. A major role of the postsynaptic density is to stabilize and anchor glutamate receptors such as AMPA and NMDA. Proteins from the Discs large (DLG) family such as Postsynaptic density protein-95 (PSD-95), also called DLG4, is involved in different protein-protein interactions in the postsynaptic density. Here, they control molecular organization and regulate synaptic strength by altering the function and distribution of AMPA receptors at the synapse (Chen et al. 2011). DLG4 contains five folded domains: Postsynaptic density-95/Discs large/Zonula occludens (PDZ)1, PDZ2, PDZ3, Src homology 3 (SH3) and guanylate kinase like (GK). DLG4 is one of four paralogs in vertebrates together with SAP97 (DLG1), PSD-93 (DLG2) and SAP102 (DLG3), respectively. These four proteins arose as a result of two consecutive whole genome duplications in the vertebrate lineage approximately 440 million years ago (McLysaght, Hokamp, and Wolfe 2002; Putnam et al. 2008). The amino acid sequences of the three PDZ domains found in each of these proteins are well conserved. In fact, the identity and similarity are high also for PDZ domains in the corresponding orthologous proteins in evolutionarily distantly related animals such as *Drosophila melanogaster* and the fresh-water polyp *Hydra vulgaris* (Fig. 1a).

**Fig 1.**
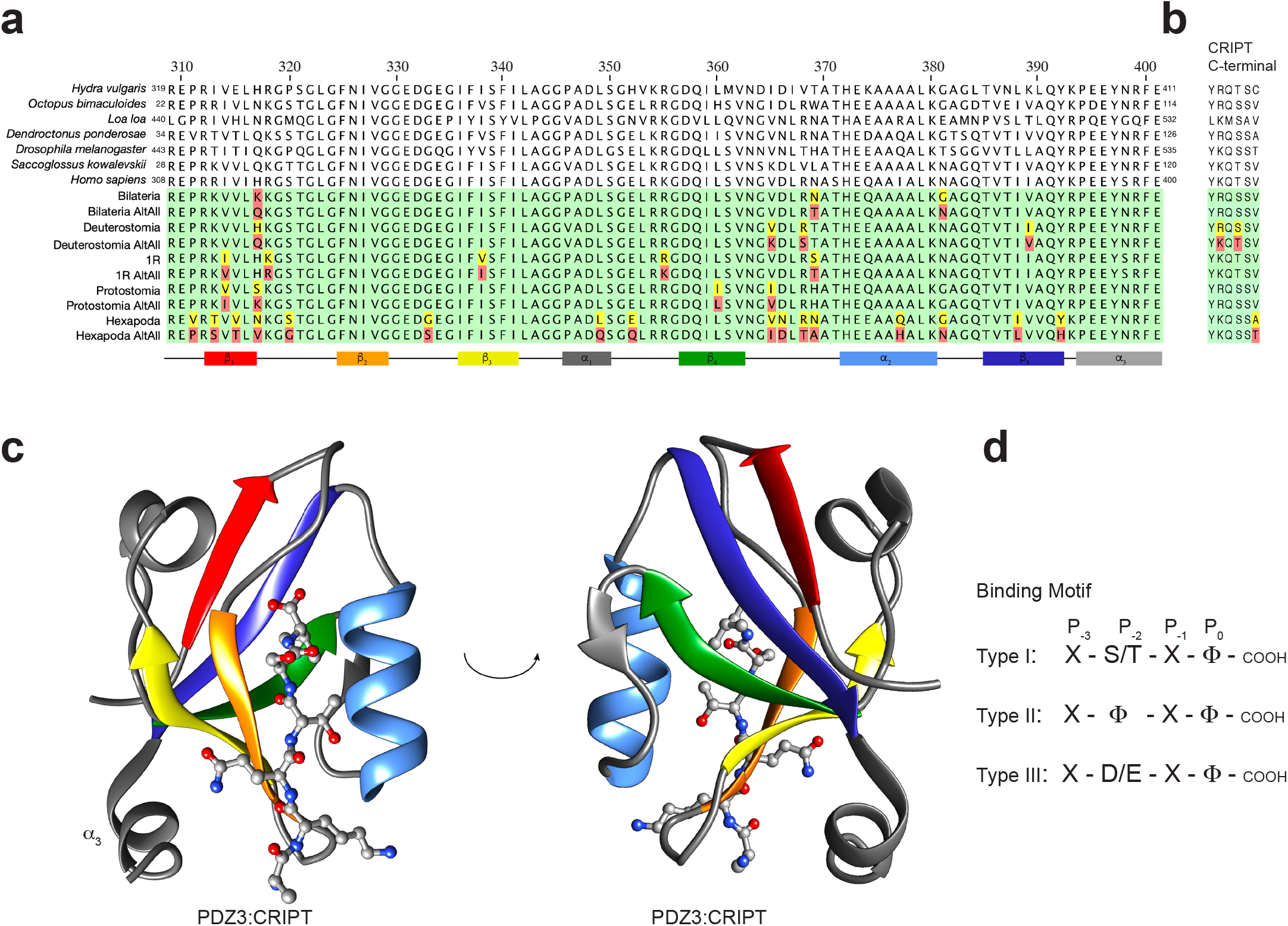
Extant and reconstructed ancient DLG PDZ3 and CRIPT sequences. **a** Alignment of DLG PDZ3 sequences. The sequence numbering of extant PDZ3 domains refers to the full-length DLG protein from the respective species. For simplicity, we use numbering of PDZ3 according to the human DLG4 sequence throughout the paper. In the alignment of ancestral sequences (green background), residues with posterior probabilities lower than 0.5 (red) and higher than 0.5 but lower than 0.8 (yellow) are highlighted. The secondary structure is from the crystal structure of DLG4 PDZ3 depicted in panel c with the corresponding color code. **b** Alignment of C-terminal CRIPT sequences. **c** Crystal structure of human DLG4 PDZ3 in complex with a CRIPT peptide (YKQTSV) (PDB ID: 5HEB). Secondary structure elements are marked in accordance with the alignment in panel a. The structure was visualised using UCSF Chimera software. **d** Illustration of three common C-terminal PDZ binding motif types. X, any amino acid residue; Φ, hydrophobic residue. Previous studies have shown that human DLG4 PDZ3 prefers a type I motif with a Val at position P_0_.

The PDZ domain is ancient and based on sequence similarity it can be traced in animals as well as in fungi and plants (Emes and Grant 2011), which contain a permuted variant (Ivarsson et al. 2008). Within the animal kingdom there are examples of genomes containing hundreds of distinct PDZ domains showing that it has been recycled as a protein-protein interaction module many times during evolution following gene or genome duplications. This abundance of PDZ domains in the animal proteome is likely facilitated by its structural architecture allowing easy integration into other proteins (Harris and Lim 2001). PDZ domains consist of 80-100 residues and have a compact globular fold usually containing 5-6 antiparallel β strands and two α helices. However, DLG PDZ3 contains a third α helix at the C-terminus (α_3_) (Fig. 1c).

Protein ligands of PDZ domains usually bind via their C-termini to a binding groove in the PDZ domain. This groove is shaped by a conserved motif in the β_1_β_2_ loop (GLGF or variants of it), the β_2_ strand and the α_2_ helix. In most cases, the C-terminus of the protein ligand arranges as an anti-parallel β strand in the groove together with the β_2_ strand (Fig. 1c). The part of the protein ligand that binds to the PDZ domain is called the PDZ binding motif (PBM). Affinity is primarily dictated by the protein ligand residues denoted P_−2_ and P_0_, where P_0_ is the C-terminal residue and P_−2_ the third residue from the C-terminus (Fig. 1d). Early studies led to classification of PBMs as type I (T/S-X-ϕ-COOH), type II (ϕ-X-ϕ-COOH) or type III (D/E-X-ϕ-COOH), based on the nature of the residues at P_−2_ and P_0_. However, the classification of PDZ domains only depending on two residues of the binding partner is an over simplification. Not only is the specificity of PDZ-ligand interactions usually dependent on more than P_−2_ and P_0_ residues of the ligand (Ernst et al. 2014), but also several properties of the PDZ domains themselves modulate binding, for example, residues outside of the binding groove (Ye et al. 2018) and extensions to the canonical PDZ domain such as α_3_ in DLG PDZ3 (Zeng et al. 2016).

One protein ligand for DLG PDZ3 is cysteine-rich interactor of PDZ3 (CRIPT) (Niethammer et al. 1998). This is an interaction that is well studied both from a structural and biophysical point of view, since it was the first structure solved of a PDZ domain with peptide ligand (Doyle 1996). Full affinity of CRIPT (*K*_d_ in the low μM range for the *Homo sapiens* CRIPT:DLG4 PDZ3 interaction) is obtained with the six last amino acid residues(Saro et al. 2007; Gianni et al. 2011; Toto et al. 2016). CRIPT has a neurological function and is required for DLG to promote dendrite growth by AMPA activation (Zhang et al. 2008; Beique et al. 2006; Zhang et al. 2017). However, certain animals including the model organisms *D. melanogaster* (where DLG was first identified) and *Caenorhabditis elegans* have CRIPT proteins that lack the classical PDZ-binding motif (Fig. 1b and Supplementary Fig. 1). In *D. melanogaster*, DLG is involved in a range of processes, including synaptic clustering of Shaker potassium channels(Tejedor et al. 1997), junction structure, cell polarity and localization of membrane proteins. Recent work shows that CRIPT clusters next to DLG in the synapse, thereby promoting dendrite growth (Zhang et al. 2017). Furthermore it was reported from knock down experiments that CRIPT is essential for DLG dependent dendrite growth (Zhang et al. 2017). Thus, there is some ambiguity whether CRIPT from, *e.g*., *D. melanogaster*, binds PDZ3 domains when it lacks the classical PDZ-binding motif and whether the biological action of DLG and CRIPT is partially independent of the PDZ3:CRIPT interaction (Zhang et al. 2017).

Evolutionary biochemistry, *i.e*., phylogenetic reconstruction of ancestral sequences followed by expression and experimental characterization of the ancient proteins (Fig. 2a-b) is a powerful tool for understanding protein function (Hochberg and Thornton 2017), including protein-protein interactions (Hultqvist et al. 2017; Jemth et al. 2018; Wheeler et al. 2018). To better understand the evolution and hence function of the PDZ3:CRIPT interaction we reconstructed sequences of ancestral time-matched variants of PDZ3 and CRIPT and compared their binding affinity and stability to PDZ3 and CRIPT from seven extant species. We show that the PDZ3:CRIPT interaction has evolved such that the affinity has been maintained or slightly increased in most extant animal phyla (chordates, hemichordates, molluscs and cnidarians) whereas affinity seems to be lost among most nematodes and insects. Our findings raise two questions: Is CRIPT the natural ligand for DLG PDZ3, and is the control of AMPA and NMDA receptors independent of PDZ3:CRIPT interaction, at least in nematodes and insects?

**Fig 2.**
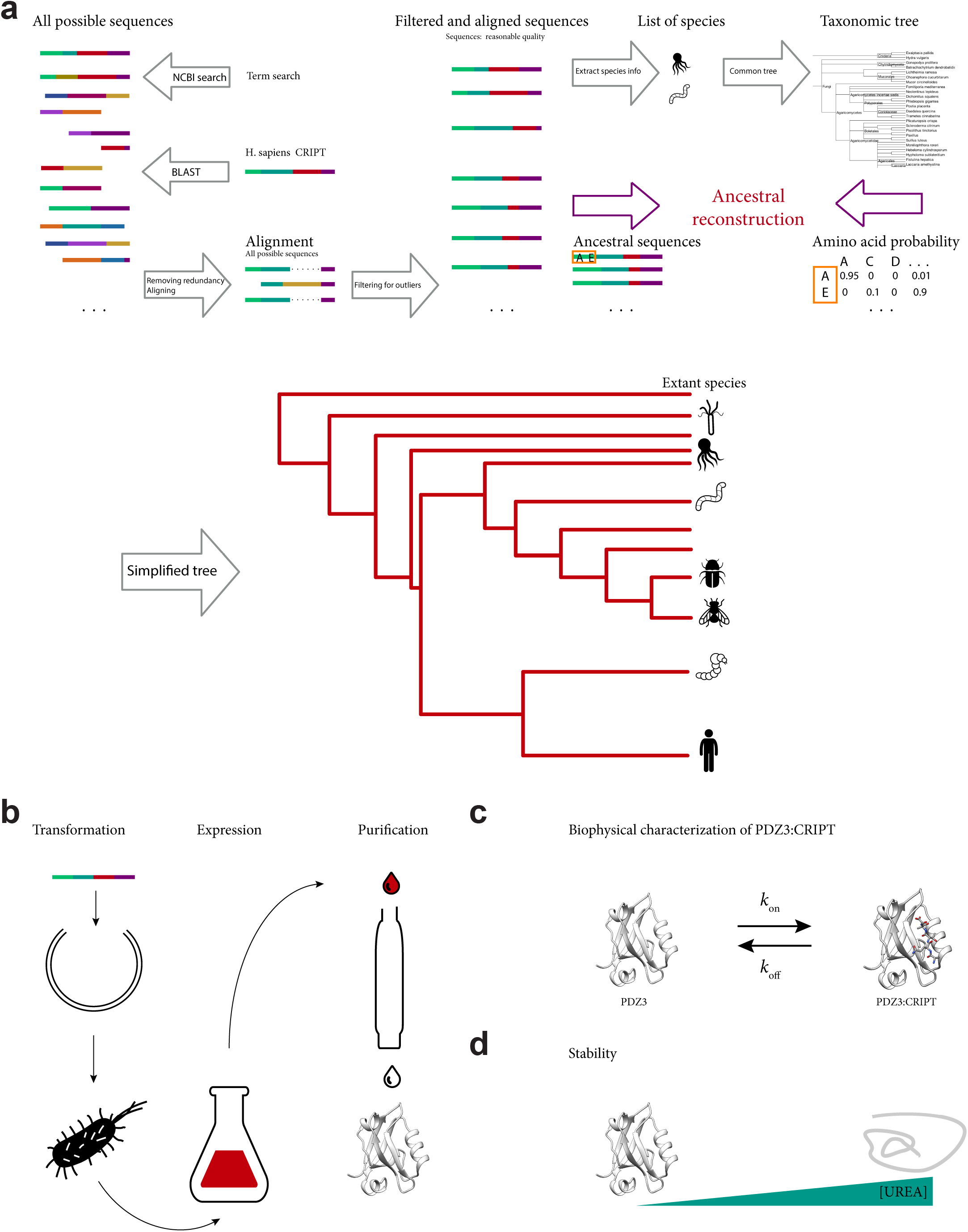
Overview of setup to investigate the evolution of the PDZ3:CRIPT interaction using evolutionary biochemistry. **a** Ancestral sequence reconstructions of CRIPT and PDZ3, respectively, were performed based on multiple sequence alignments and a species tree. In the reconstruction, 309 unique sequences of CRIPT from 498 different species and 249 unique sequences of PDZ3 from the DLG protein family from 324 different species were included. **b** Expression and purification of selected extant and reconstructed ancestral protein variants. **c** The PDZ3:CRIPT interaction was analyzed by isothermal titration calorimetry and stopped-flow spectroscopy. **d** Global stability of PDZ3 domains were determined by circular dichroism-monitored urea denaturation experiments.

## Results

### Reconstruction of ancestral sequences

Protein sequences for DLG family me mbers (including DLG4 and its vertebrate paralogs DLG1, DLG2 and DLG3) and CRIPT proteins were recovered from the NCBI database (NCBI Resource Coordinators). The final set of sequences for ancestral reconstruction consisted of 309 unique sequences of CRIPT from 498 different species and 249 unique sequences of DLG family PDZ3 domains from 324 different species (Supplementary Fig. 1 and 2). The number of different species exceeded the number of unique sequences due to 100% sequence identity between several closely related species. While all of the PDZ3 sequences were from Metazoan (animal) species, CRIPT sequences also included other kingdoms since homologous proteins with high sequence similarity were found in both fungi and plants. Within the animal kingdom, the similarity between homologous sequences of both DLG PDZ3 and CRIPT was very high (Supplementary Fig. 1 and 2). Availability and quality of sequences were variable so the widest most representative dataset was constructed using search methods, sequence alignment, filtering and sorting using Python programming and manual curation. Due to high sequence similarity, posterior probabilities for the maximum likelihood estimate (ML) of reconstructed ancestral sequences were relatively high for most positions in the alignment (Supplementary Data Excel file 1 and 2). We were able to reconstruct PDZ3 from five historical time points, namely from the common ancestor of all extant bilaterians, deuterostomes, protostomes, and hexapods (insects), respectively, as well as from the common ancestor of all extant vertebrates containing four DLG paralogs (Fig. 1a). The PDZ3 from the latter ancestor is denoted 1R PDZ3, since the four DLG paralogs arose as a result of two consecutive whole genome duplications called 1R and 2R, respectively (McLysaght, Hokamp, and Wolfe 2002; Putnam et al. 2008). The C-terminus, *i.e*., the PBM of CRIPT was reconstructed at eight historical time points, namely from the common ancestors of all extant eukaryotes, opisthokonts (animals and fungi), eumetazoans (animals except sponges), bilaterians, deuterostomes, protostomes, vertebrates (1R) and hexapods, respectively (Fig. 1b and Supplementary Fig. 3). In addition to the ancient ones, we expressed and characterized DLG PDZ3 and CRIPT variants from the following extant species representing different distantly related animal groups: the non-bilaterian animal *Hydra vulgaris* (a cnidarian), *Octopus bimaculoides* (California two-spot octopus, a mollusc and protostome), *Loa loa* ("eye worm", a nematode or roundworm causing the disease loiasis, protostome), *Dendroctonus ponderosae* (mountain pine beetle, an insect, protostome), *Drosophila melanogaster* (fruit fly, insect, protostome), *Saccoglossus kowalevskii* (acorn worm, a hemichordate, deuterostome) and *Homo sapiens* (representing chordates, deuterostome) (Fig. 2 and Supplementary Fig. 3). We use “native interaction” to denote interactions between time- and species matched PDZ3:CRIPT interactions. Based on sequence alignment and structure prediction, it appears that all ancestral and extant DLG PDZ3 domains have a similar structure with the usual PDZ fold consisting of six β strands and two α helices, but also including a third C-terminal α helix (α_3_), present in the crystal structures of human DLG4 PDZ3 (Doyle et al. 1996) (Supplementary Fig. 4).

### Evolution of PDZ3:CRIPT interaction

From the sequence alignment and phylogeny of the C-terminal residues in CRIPT (Supplementary Fig. 1) we can follow the evolution of the PBM (Supplementary Fig. 3). Among viridiplantae, plants and mosses show a striking and complete conservation of a type 1 motif (YKQSNV), while green algae have a polar residue at position P_0_. The majority of animal phyla including vertebrates, annelids and molluscs (except Euthyneura, i.e., snails and slugs) have CRIPT with a classical type 1 PBM. However, fungi and arthropods (including insects, spiders and crustaceans) appear to lack a PBM, although there are a few exceptions. Interestingly, CRIPT from the cnidarian *H. vulgaris* ends with a Cys (Fig. 1b).

In accordance with the alignment and phylogeny, CRIPT from the last common ancestor of all extant Eukaryotes contained a classical type 1 PBM (YKQSSV) (Supplementary Fig. 3). A type 1 PBM was likely also present in the ancestor of all animals but has since mutated in distinct animal phyla and orders. Clearly, the ancestor of bilaterian animals contained a type 1 PBM and so did the ancestors of the two major bilaterian groups, protostomes and deuterostomes. However, the last common ancestor of today’s arthropods likely carried a mutated PBM in CRIPT, ending with Ala or Thr, instead of the canonical Val residue.

One intriguing notion is that the C-terminal residues P_−1_ to P_−5_ are highly conserved even among CRIPTs with Ala or Thr at position P_0_. This raises the question whether CRIPTs without a canonical type 1 PBM still retain binding to their native DLG PDZ3? To address this question, PDZ3 and CRIPT from seven extant and five ancestral species were subjected to binding experiments using ITC and stopped-flow spectroscopy (Fig. 3 and 4).

**Fig 3.**
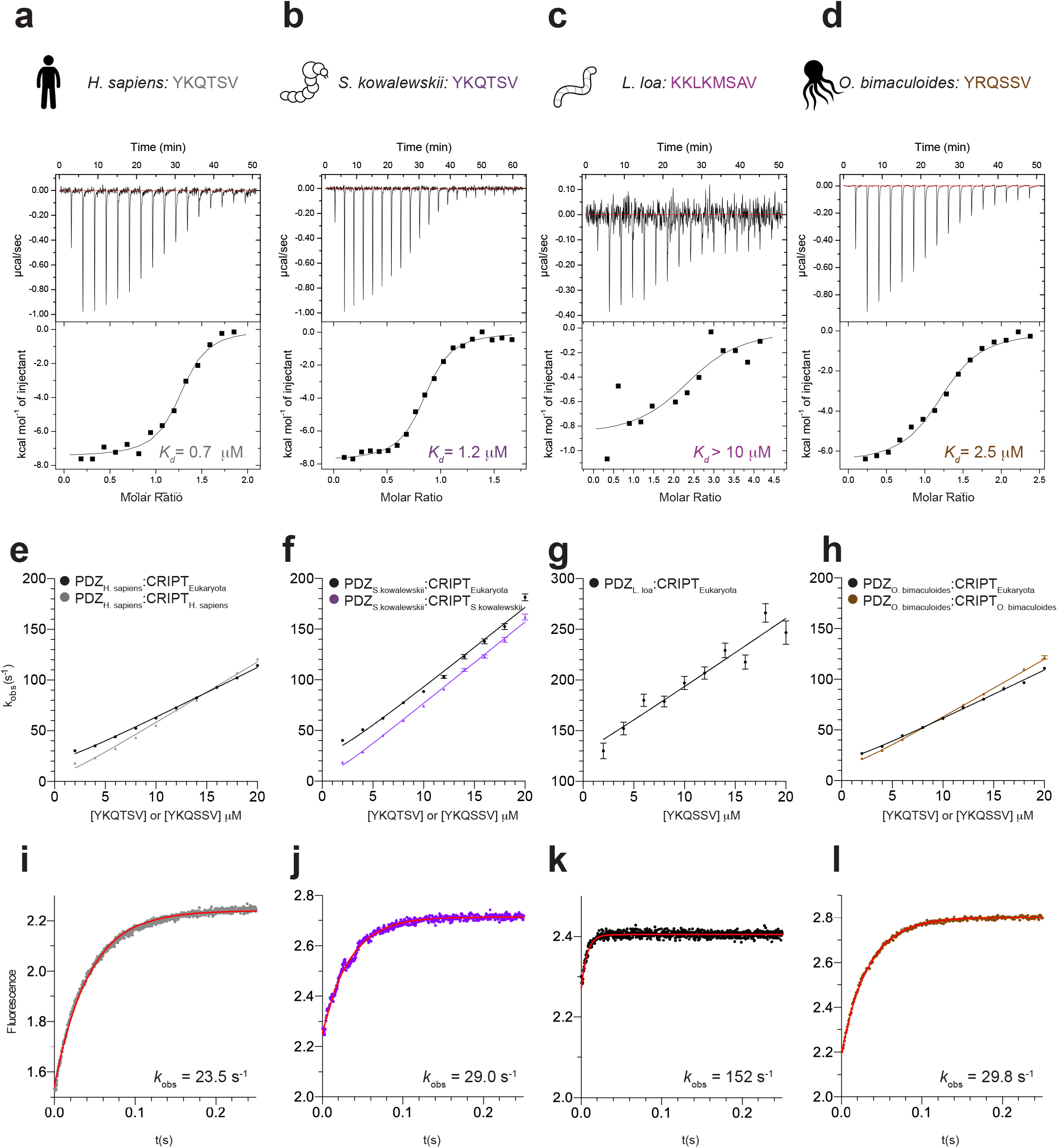
Characterization of the PDZ3:CRIPT interaction from extant species with a type 1 PBM in CRIPT. **a-d** Examples of ITC experiments for native PDZ3:CRIPT interactions for four extant species: *H. sapiens*, *S. kowalewski*, *L. loa* and *O. bimaculoides*. **e-l** Binding kinetics from stopped-flow experiments for the same native interactions as in panel **a-d**, and also including experiments with a reconstructed CRIPT from the ancestor of all eukaryotes (CRIPT_Eukaryota_). **e-h** Observed rate constants were obtained from kinetic binding traces (examples shown in panels **i-l**) and plotted as function of CRIPT concentration at a constant concentration of PDZ3 (1 μM) to estimate the association rate constant *k*_on_. **i-l** Examples of kinetic binding traces from stopped-flow experiments using 4 μM CRIPT and 1 μM PDZ3. The fitted line (red) is a single exponential from which the observed rate constant *k*_obs_ was obtained. *k*_off_ was determined in a separate experiment as explained in the Methods section, and *K*_d_ calculated as *k*_off_/*k*_on_. Experiments were performed at 25°C for ITC and at 10°C for stopped-flow.

**Fig 4.**
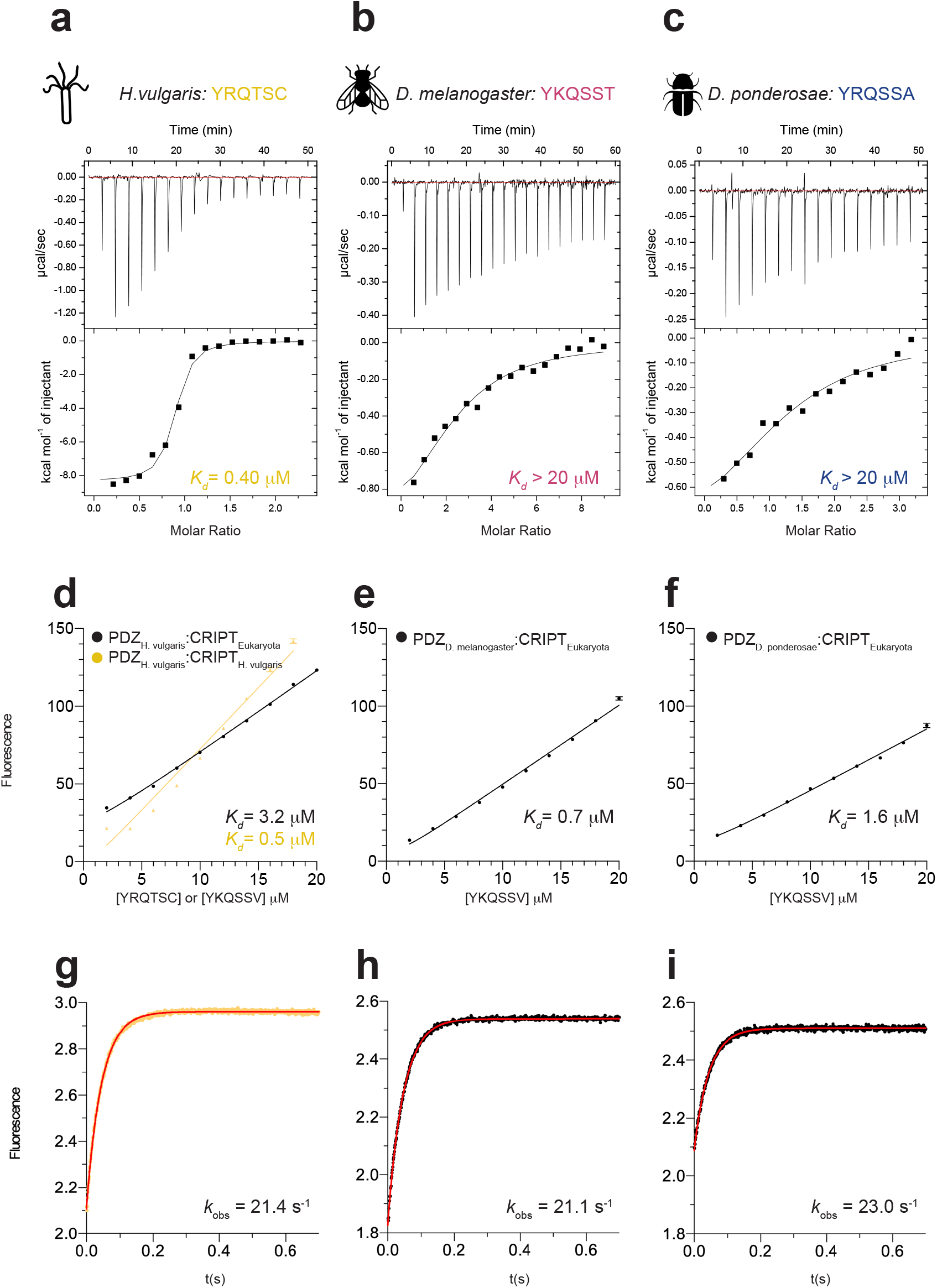
Characterization of the PDZ3:CRIPT interaction from extant species without a type 1 PBM in CRIPT. **a-c** Examples of ITC experiments for native PDZ3:CRIPT interactions for three extant species (*H. vulgaris*, *D. melanogaster* and *D. ponderosae*, respectively). **d-i** Binding kinetics from stopped-flow experiments for the native *H. vulgaris* interaction and also experiments with the respective PDZ3 and a reconstructed CRIPT from the ancestor of all eukaryotes (CRIPT_Eukaryota_). **d-f** Observed rate constants were obtained from kinetic binding traces (examples shown in panels **g-i**) and plotted as function of CRIPT concentration at a constant concentration of PDZ3 (1 μM) to estimate the association rate constant *k*_on_. **g-i** Examples of kinetic binding traces from stopped-flow experiments using 4 μM CRIPT and 1 μM PDZ3. The fitted line (red) is a single exponential from which the observed rate constant *k*_obs_ was obtained. *k*_off_ was determined in a separate experiment as explained in the Methods section, and *K*_d_ calculated as *k*_off_/*k*_on_. Experiments were performed at 25°C for ITC and at 10°C for stopped-flow.

The binding of *H. sapiens* CRIPT to *H. sapiens* DLG4 PDZ3 is well characterized (Niethammer et al. 1998; Saro et al. 2007; Gianni et al. 2011; Toto et al. 2016; Doyle et al. 1996), and we report a similar binding affinity (0.4 μM) as previous studies (Fig. 3 and 5, Table 1 and 2). Furthermore, PDZ3 from *H. vulgaris* (with a C-terminal Cys residue), *O. bimaculoides, S. kowalevskii* and the common ancestors of bilaterial animals, protostomes, deuterostomes, and vertebrates (1R), respectively, were all found to bind to their time-matched native CRIPT with low μM affinity (Fig. 3, 4 and 5, Table 1 and 2). On the other hand, PDZ3 domains from *D. melanogaster*, *D. ponderosae* and the ancestor of hexapods bind poorly to their respective native CRIPT, which lacks a type 1 PBM. While the interactions were too weak for stopped-flow spectroscopy, ITC experiments provided a rough estimate of the affinities (*K*_d_ ≥ 10-30 μM) (Table 2, Fig. 4 and 5). More surprisingly, PDZ3 from the nematode *L. loa* was also found to bind poorly to its native CRIPT even though it contains a type 1 PBM with a C-terminal Val (Fig. 3).

**Fig 5.**
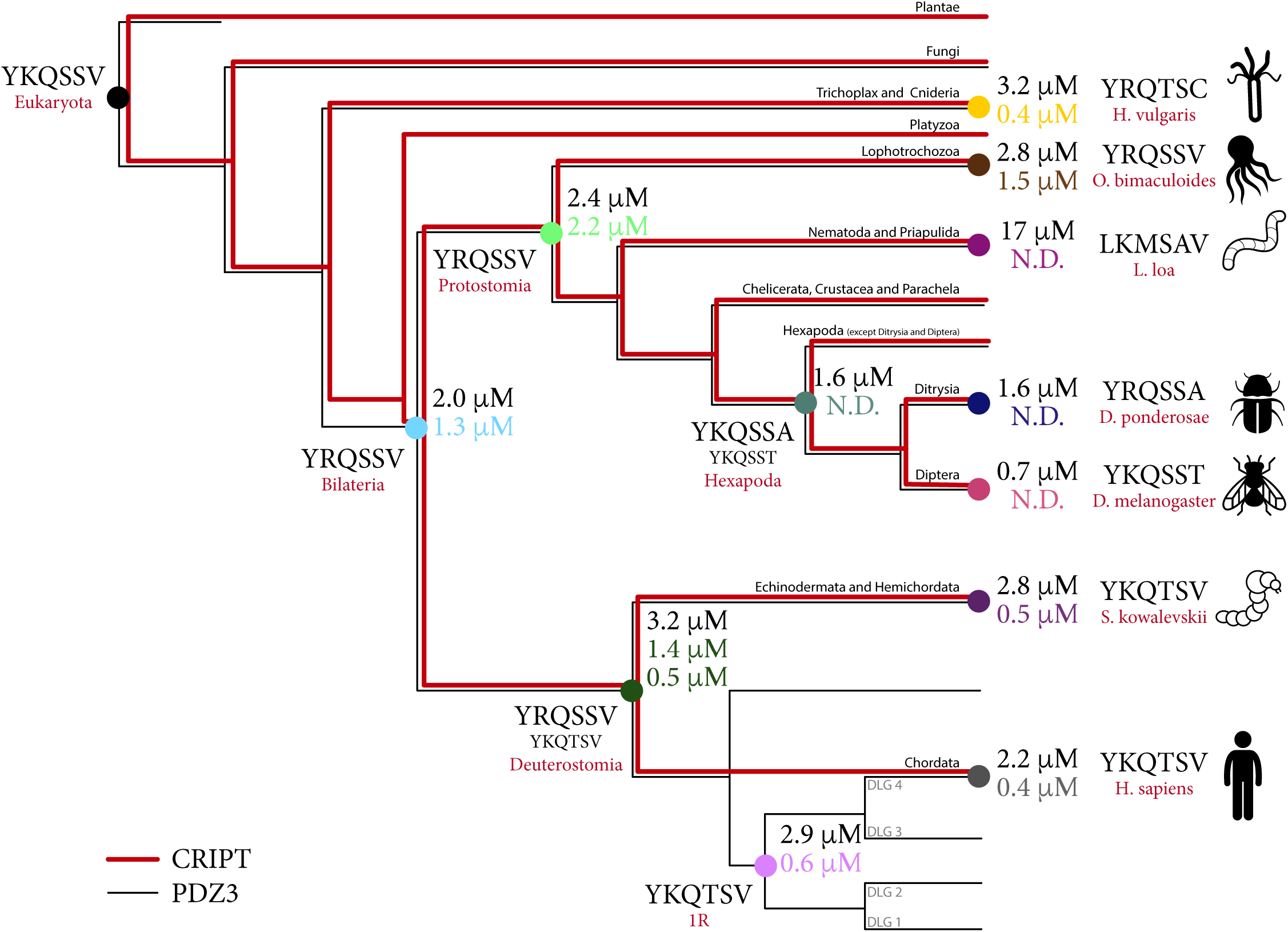
Evolution of affinity for the PDZ3:CRIPT interaction. Simplified species tree with nodes for selected ancestral and extant species used in the study. The native time-matched affinity for each interaction is shown in colour code text at each node and the affinity between each PDZ3 and Eukaryota CRIPT is shown in black text. For the native interaction in the deuterostome node, two CRIPT peptides were tested due to low posterior probability for Ser at position P_−2_. *K_d_* values were determined by stopped-flow experiments at 10°C.

**Table 1.**
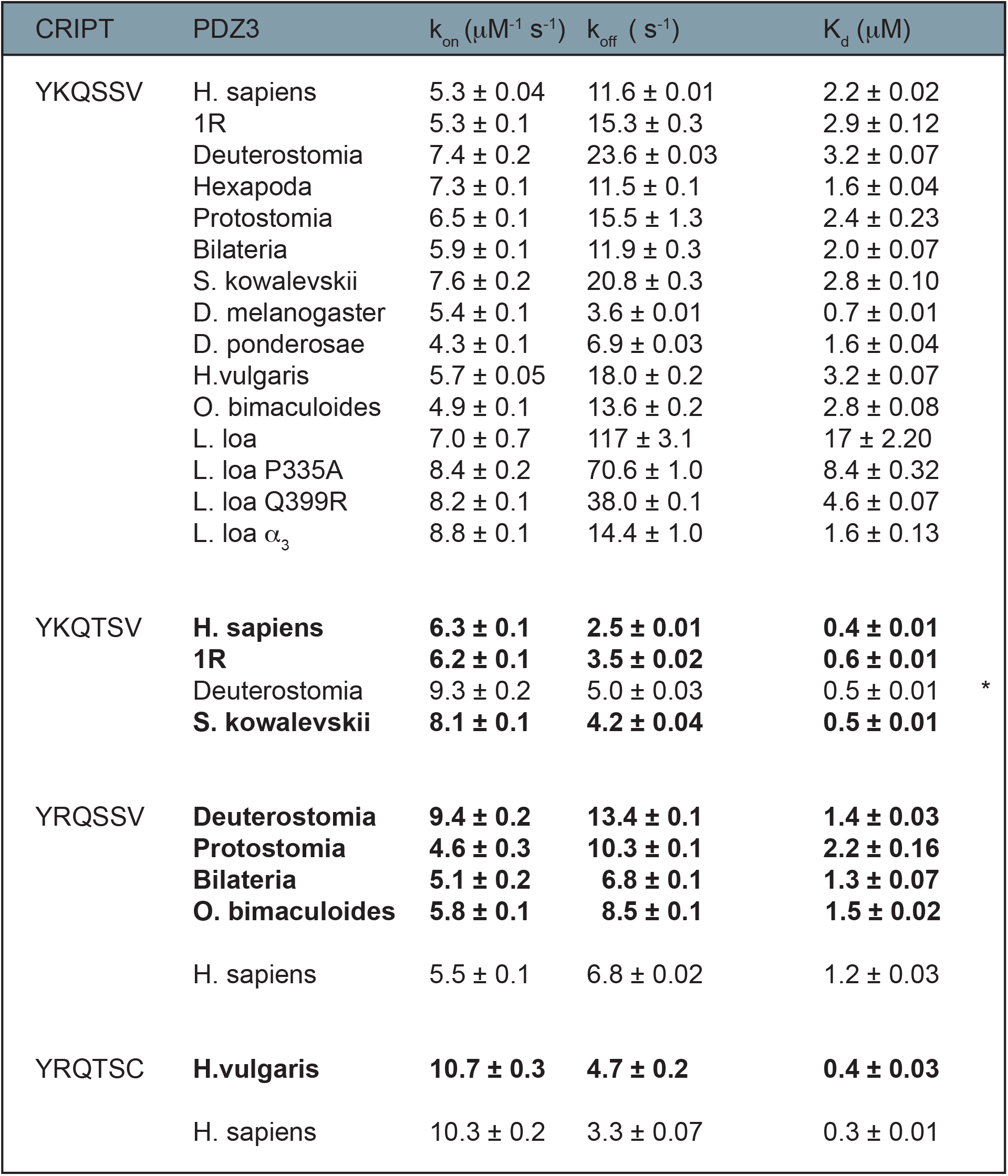
Binding constants for extant and reconstructed ancestral PDZ3:CRIPT interactions from kinetic experiments. Rate and equilibrium constants were determined for the PDZ3:CRIPT interaction for extant and ancestral PDZ3 domains binding its respective native CRIPT (YKQTSV, YRQSSV or YRQTSC). In addition, binding experiments were performed for all PDZ3 domains to the ancestral Eukaryota CRIPT (YKQSSV) and for all CRIPTs to *H. sapiens* PDZ3. Native interactions are highlighted in bold. Stopped-flow experiments were used to determine the association rate constant (*k*_*on*_) from binding experiments and the dissociation rate constant (*k*_*off*_) from displacement experiments. *K*_*d*_ values were calculated from the ratio of *k*_*off*_ and *k*_*on*_. All data were obtained at 10°C in 50 mM sodium phosphate, pH 7.45, 21 mM KCl (I = 150).

**Table 2.**
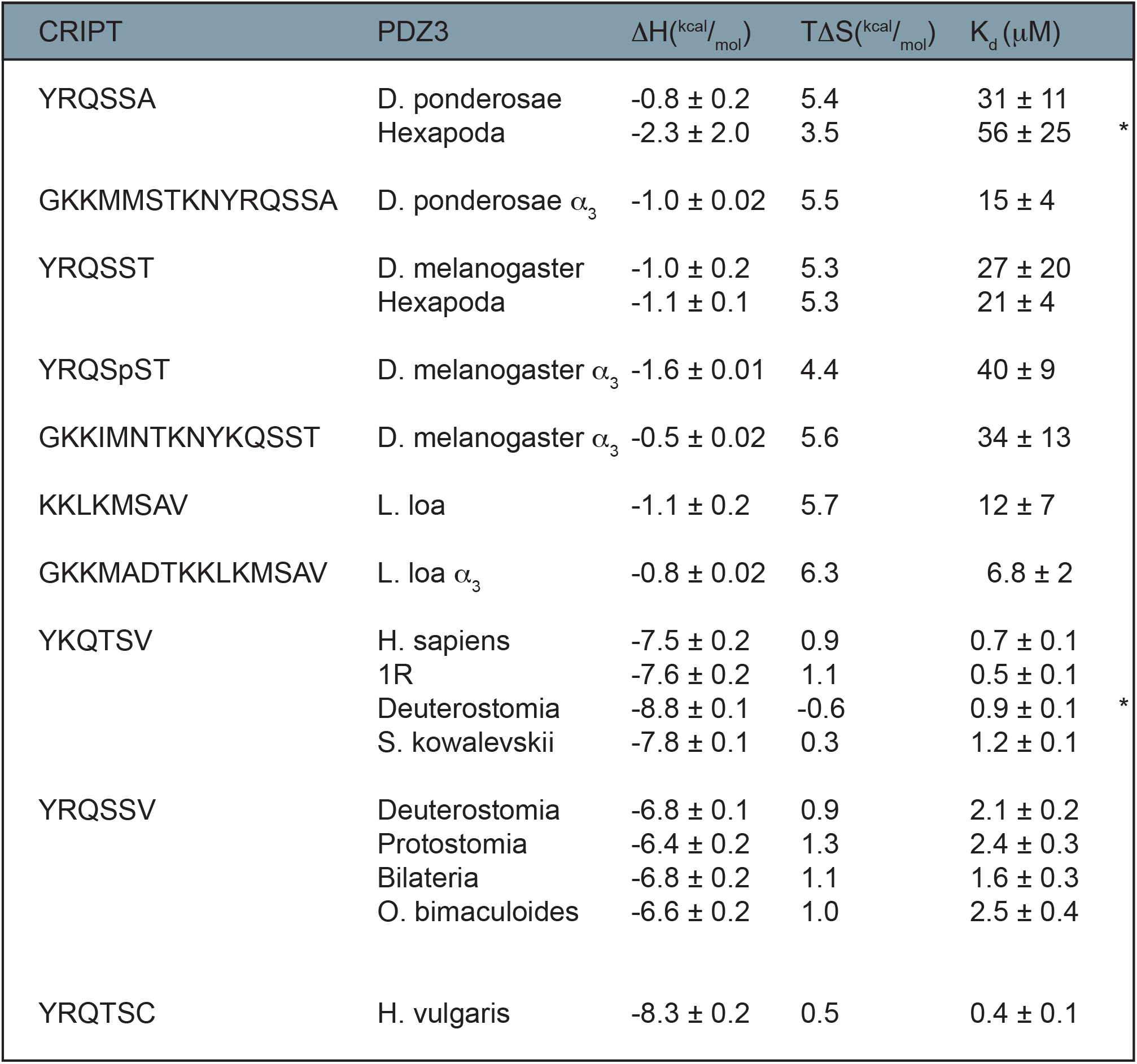
Binding constants for extant and reconstructed ancestral PDZ3:CRIPT interactions monitored by ITC. Thermodynamic parameters for native ML or AltAll(*) PDZ:CRIPT interactions were determined at 25°C in 50 mM sodium phosphate, pH 7.45, 21 mM KCl (I = 150). A longer *L. loa* CRIPT peptide was used (8 residues) due to low solubility of the hexamer.

To directly compare each PDZ3 domain with regard to binding of CRIPT with a type 1 PBM, the affinity of CRIPT from the ancestor of all eukaryotes (YKQSSV) was measured with all PDZ3 domains included in the study. The ancestral Eukaryota CRIPT was found to bind to all resurrected and present-day PDZ3 domains, including insect PDZ3 domains, in the low μM range (Fig 3, 4 and 5, Table 1). However, the binding of Eukaryota CRIPT to the nematode *L. loa* PDZ3 was somewhat weaker than for other PDZ3 domains (17 μM), suggesting that *L. loa* PDZ3 has a binding groove with a slightly different character than PDZ3 domains from the other investigated extant and ancestral species. Thus, while nematodes, the sister phylum of arthropods, has retained the type 1 PBM in CRIPT the high affinity of the PDZ3:CRIPT interaction is lost, suggesting that it may not be functional in either group of animals. Another option is that the PDZ3:CRIPT interaction is functional but interactions outside the canonical groove is essential for full affinity among Ecdysozoa as observed for some other PDZ:ligand complexes (Ye, Huang et al. 2018) (Zeng et al. 2016; Li et al. 2014; Pan et al. 2011) or that the PDZ3:CRIPT interaction is posttranslationally regulated (Sundell et al. 2018). Obviously, lower affinity could have evolved as a response to a certain course of events leading to for example high local cellular concentrations of DLG and CRIPT. Finally, a trivial explanation is that the PDZ3 domains used in the experiments were not properly folded. While it is challenging to test intracellular expression levels we attempted to partially address the other issues.

### Mutational analysis to resolve the low CRIPT affinity of PDZ3 from *L. loa*

The *L. loa* PDZ3:CRIPT interaction is particularly interesting since it appears as if the PDZ3 domain has lost affinity for CRIPT with type 1 PBM. Sequence alignment (Fig. 1a) and structural prediction of the PDZ3 variants show that *L. loa* PDZ3 differs in several amino acid positions as compared to other PDZ3 domains, which inspired us to check whether interactions outside the canonical groove could be essential for the PDZ3:CRIPT interaction. *L. loa* PDZ3 has several unique residues at positions that previously have been reported to be important for binding affinity (Supplementary Fig. 5), *e.g*. , Gly→Pro at position 335 in the β_2_β_3_ loop next to the canonical binding groove and Arg→ Gln at position 399 in α_3._ The presence of Pro335 and/or Gln399 is rare; they are only found in 12 sequences among the 253 sequences used for the ancestral sequence reconstruction. Furthermore, a previous study showed the significance of the β_2_β_3_ loop for increasing affinity by stabilization of a salt-bridge formed by Arg 399 in α_3_ and Glu 334 in the β_2_β_3_ loop in *H. sapiens* DLG4 PDZ3 (Mostarda, Gfeller, and Rao 2012). Thus, two single point mutations, P335G and Q399R, were introduced into *L. loa* PDZ3, to test if the amino acid substitutions can explain the weak binding to CRIPT. Indeed, the affinity increased two and three-fold, respectively, towards ancestral Eukaryota CRIPT (Table 1), supporting the notion that the derived substitutions in *L. loa* PDZ3 indeed weakened the interaction with CRIPT.

### Global stability of ancestral and extant PDZ3 domains

Reconstructed ancestral proteins often show a higher stability than the present-day proteins they were predicted from. One biological explanation (for ectotherm organisms) is the higher environmental temperatures in the past (Gaucher, Govindarajan, and Ganesh 2008; Risso, Gavira, and Sanchez-Ruiz 2014), but others argue that the higher stability of ancient proteins is an artifact of the sequence reconstruction because of a consensus sequence bias in the approach(Trudeau, Kaltenbach, and Tawfik 2016; Sternke, Tripp, and Barrick 2019; Williams et al. 2006).

We determined the global stability of all selected extant and resurrected ancient PDZ3 variants using urea denaturation experiments monitored by far UV circular dichroism, which detects secondary structure and is a robust probe of global folding (Fig. 6a-d, Table 3). It is clear that all PDZ3 domains in the study are well folded under our experimental conditions and that they unfold by an apparent two state mechanism in equilibrium urea denaturation experiments. Thus, our binding data are not influenced by low stability of ancient or extant proteins. However, because of uncertainties in curve fitting, as described below, we were careful in the interpretation. All resurrected PDZ3 domains were found to be more stable in comparison to extant PDZ3 domains except *H. sapiens* PDZ3. Generally, we observe a slight increase in the urea midpoint for the deuterostome lineage from bilaterian to *H. sapiens* DLG4 PDZ3. However, there is some ambiguity in the actual stability of the PDZ3 variants due to uncertainty in determination of the *m*_D-N_ value, which reflects the change in solvent accessible hydrophobic surface area upon denaturation and which is directly related to the stability: Δ*G*_*D-N*_ = [Urea]_50%_ × *m*_D-N_. It is fair to assume that all PDZ3 variants of similar sequence length also have a similar *m*_D-N_ value. Therefore, we fitted the data with a shared *m*_D-N_ value, which gives a more robust estimate of the differences in Δ*G*_*D-N*_ between PDZ3 variants. But we also fitted the data with a free *m*_D-N_ value to get an idea of the error in this parameter. In addition, some PDZ3 variants did not show complete unfolding at the highest urea concentration, resulting in a lower accuracy of the fitted parameters. The described uncertainties are not unique to our study but pertain to all studies where thermodynamic stabilities are compared, including previous studies addressing ancient versus modern proteins.

**Fig 6.**
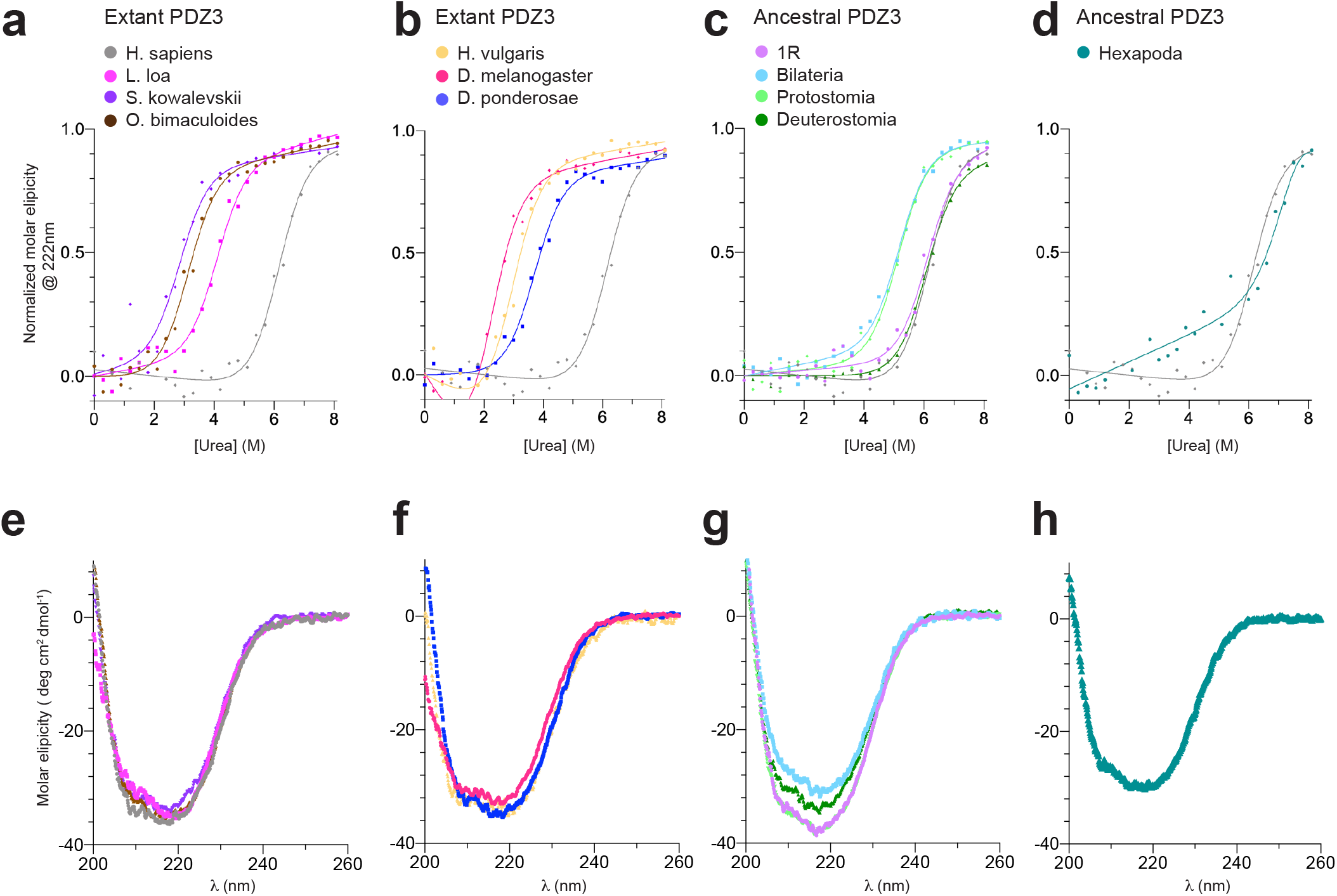
Secondary structure and global stability of extant and resurrected ancestral PDZ3 domains. **a-d** Urea denaturation (0-8.1 M) of extant and ancestral PDZ3 variants, as monitored by circular dichroism at 222 nm. Data were fitted to a two-state model for protein (un)folding. See Table 3 for fitted parameters. To facilitate comparison, the urea denaturation curve of *H. sapiens* PDZ3 (grey) is present in all graphs. **e-h**) Secondary structure content analysed by circular dichroism between 200-260 nm. Each spectrum is an average of 5 individual scans measured at 10°C in 50 mM sodium phosphate, pH 7.45, 21 mM KCl (I = 150 mM).

**Table 3.**
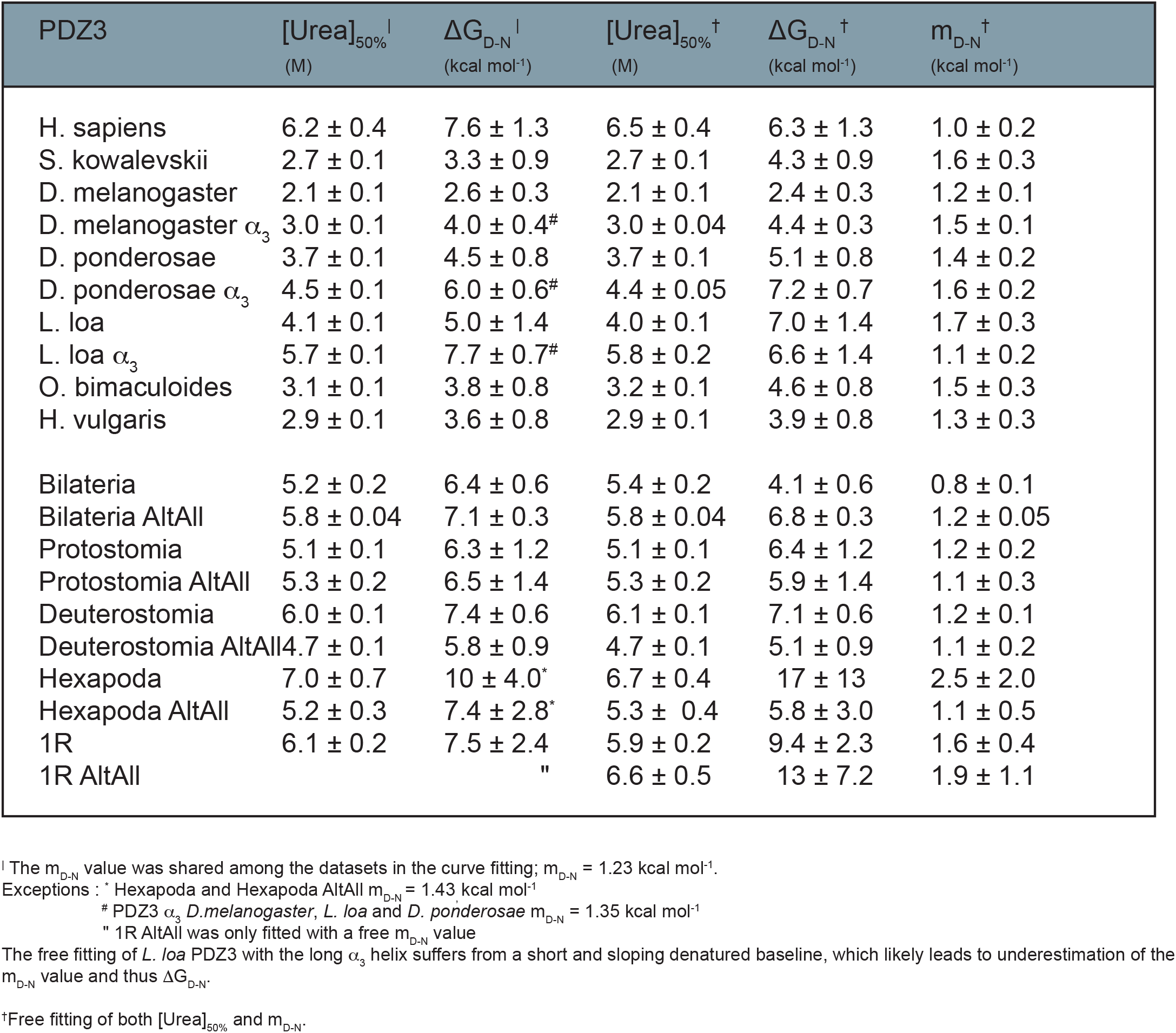
Global stability of PDZ3 domains. Global stability of PDZ3 domains was determined by urea denaturation experiments monitored by circular dichroism at 222 nm (see Fig. 6). Experimental data were fitted to a two state model for denaturation to obtain the concentration of urea where 50% of the protein is denatured ([Urea]_50%_), the cooperativity of the unfolding (m_D-N_ value) and the stability (ΔG_D-N_) as the product of [Urea]_50%_.and m_D-N_.

### The α_3_ extension increases the stability of PDZ3

Most studies of DLG4 PDZ3 have been performed on a construct based on the original crystal structure (Doyle 1996). However, in DLG4, PDZ3 is part of the supramodule PDZ3-SH3-GK in which an extended α helix (α_3_) connects PDZ3 and SH3 (Fig. 7a), but only 7 of the 19 amino acid residues of the linker are present in the constructs investigated in the present work and in previous publications. Recent studies have reported that the entire α_3_ and a peptide ligand longer than 6 residues are required for specific PDZ:ligand interactions (Zeng et al. 2016) (Zeng et al. 2018) (Ye et al. 2018). The α_3_ extension does not significantly affect the affinity between *H. sapiens* PDZ3-SH3-GK and a 6-mer or 15-mer CRIPT peptide (Laursen et al. 2019). However, as compared to the ancestral bilaterian PDZ3, the α_3_ region is highly conserved in *H. sapiens* PDZ3 with only two substitutions, whereas *D. ponderosae*, *D. melanogaster* and *L. loa* PDZ3 have 4, 8 and 9 substitutions, respectively, in the 19 residue long primary structure of α_3_ (Fig 7a). We calculated the α helix propensity of isolated α_3_ (residues 395-413) for all PDZ3 variants in the present study using the AGADIR software (Munoz and Serrano 1994). The helical propensity varied from 2-17 % (Fig. 7a), suggesting that the stability of α_3_ may have changed during evolution. (The helical propensity refers to an isolated peptide but it could correlate with stability in the context of the folded protein.) Interestingly, *L. loa*, *D. melanogaster* and *D. ponderosae* PDZ3 have the highest helix propensity in α_3_. Therefore, we decided to express, purify and analyze PDZ3 from these three species with a full length 19 residue α_3_ to assess the effect on binding and stability. We observed a slightly higher thermodynamic stability for the longer variants of PDZ3 in comparison to PDZ3 domains without an extended α_3_ (Fig 7b-d and Table 3). Similarly to the PDZ3 variants without α_3_ extension, the affinities between the extended α_3_ PDZ3 variants and their respective native CRIPT ligands (15-mer) were too low to be measured by stopped-flow. But with ITC it was again possible to obtain a rough estimate of the *K*_d_ values. The affinity between *D. melanogaster* PDZ3 and CRIPT was not affected by either the α_3_ extension or a longer CRIPT peptide (Table 2). Similarly, while the affinity seems to increase 2-fold for the respective native *D. ponderosae* and *L. loa* PDZ3:CRIPT interactions, this effect is within error of the experiment (Table 2). However, the α_3_ extension increased the affinity of *L. loa* PDZ3 for the Eukaryota CRIPT significantly, by 10-fold (Table 1), suggesting that the α_3_ extension of PDZ3, perhaps even in the context of a supramodule, needs to be considered if a weak interaction is analyzed.

**Fig 7.**
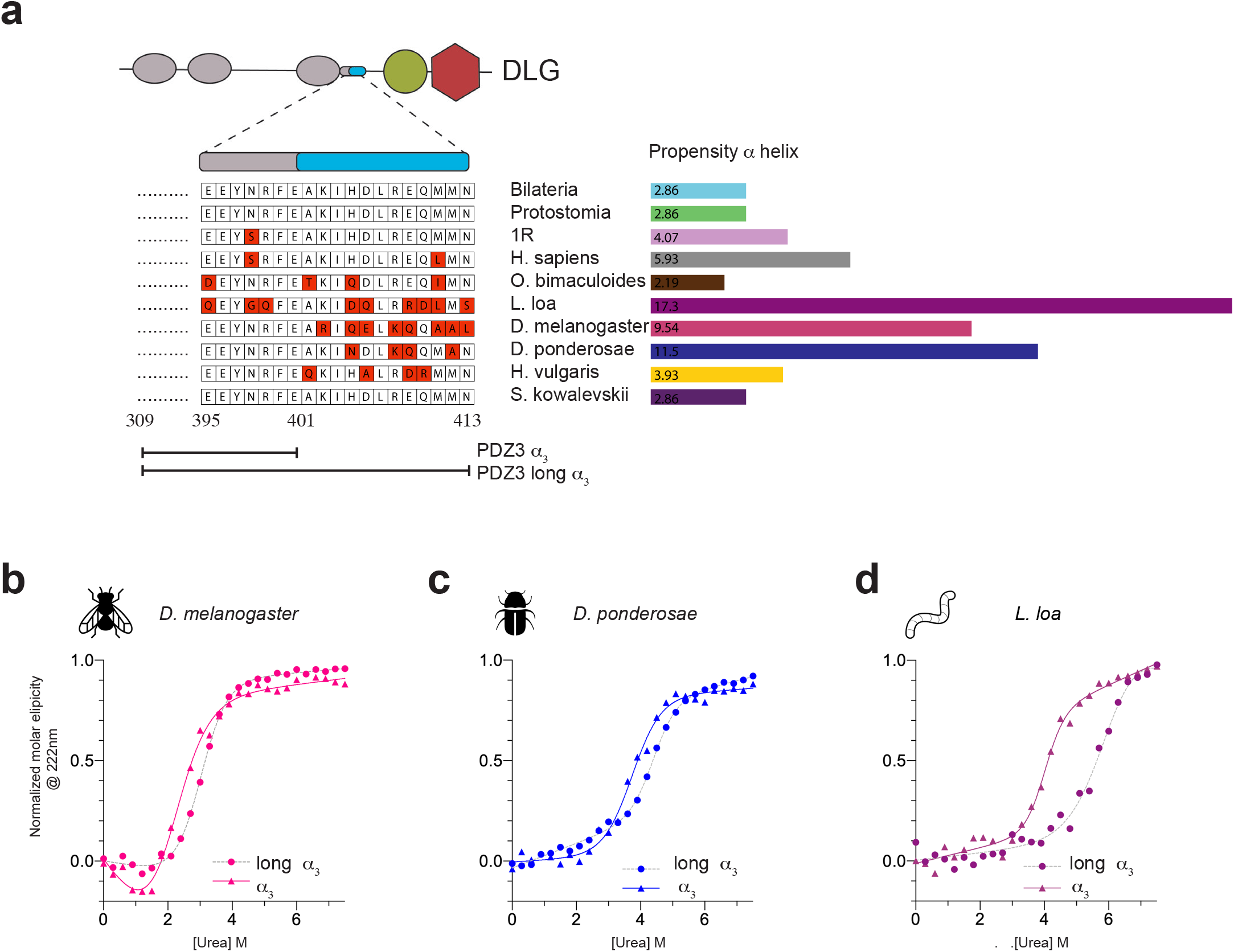
Properties of PDZ3 domains with an extended α_3_. **a** Schematic illustration of DLG4 with the PDZ1, PDZ2, PDZ3, SH3 and GK domains together with a sequence alignment of the extended α_3_ helix (19 aa) for the PDZ3 variants from extant and ancestral species included in the study. Alpha helix propensity was calculated by the AGADIR software at 283K, *I* = 150 mM, pH 7.45. **b-d** Urea denaturation experiments of PDZ3 from *D. melanogaster*, *L. loa* and *D. ponderosae*, respectively, with and without the α_3_ extension, as monitored by circular dichroism at 222 nm. See Table 3 for fitted parameters.

### Negative regulation by phosphorylation

Reversible phosphorylation results in positive or negative regulation of protein-protein interactions and therefore plays an important role for cellular signaling. Protein phosphorylation mainly occurs at Ser, Thr and Tyr residues. Thus, with regard to the PDZ3:CRIPT interaction, phosphorylation at P_−2_ (Thr) in CRIPT disables binding to PDZ3 (Sundell et al. 2018). However *D. melanogaster* CRIPT lacks the type 1 motif but has three possible phosphorylation sites, ending with SST. Therefore we tested the hypothesis that the weak interaction between *D. melanogaster* PDZ3 and its native CRIPT is positively regulated by phosphorylation of CRIPT. Phosphorylation of *D. melanogaster* CRIPT at Ser_−1_ did not affect the binding (Table 2), whereas phosphorylation of Ser_−2_ and Thr_0_ seem to weaken the affinity further since no binding could be detected by either stopped-flow or ITC. Thus, if anything, the PDZ3:CRIPT interaction is negatively regulated by phosphorylation in *D. melanogaster*.

### The conclusions based on resurrected maximum likelihood variants of PDZ3 are robust to sequence uncertainty

Finally, our historical description of the PDZ3:CRIPT evolution is contingent on accurate native affinities and stabilities for the resurrected proteins. Thus, even if the reconstructed sequences are not correct, what about the experimantally determined properties of the proteins? Different approaches can be applied to check the robustness of results based on the ML variant, for example generating a library of variants of the ML ancestral sequence by point mutations (Yokoyama and Radlwimmer 2001) or using Bayesian sampling to reconstruct a set of sequences by sampling amino acids over the posterior probability distribution (Yang and Rannala 1997). However an easier approach is to reconstruct a "worst-case scenario" variant. Here, one sequence is reconstructed in which every amino acid residue with a posterior probability below a certain threshold, for example 0.8, is replaced by the residue with the second highest probability at that position. Such a variant, denoted AltAll (alternative residue at all positions), represents a very unlikely worst case scenario that can be used to assess how robust conclusions are to errors in the reconstructed ML sequences (Eick et al. 2017; Anderson et al. 2016) simply by comparing the ML and AltAll proteins. If they display the same affinity for a ligand, that affinity is very likely to be the true ancestral affinity (Fig. 1a and Supplementary Fig. 3 and 6).

Thus, we used the AltAll approach as a relatively strong test of robustness with a single protein variant as control. Worst-case scenario AltAll sequences from CRIPT and PDZ3 were reconstructed, expressed and subjected to the same experiments as the ML variants to validate the conclusions based on the resurrected ML proteins. The C-termini of ancestral CRIPT could be reconstructed with high confidence due to the strong conservation observed in extant species. Therefore, AltAll CRIPT variants were only tested in two cases, for the ancestor of hexapods and ancestor of deuterostomes, respectively, both of which had one ambiguous position (Supplementary Fig. 3 and 6). AltAll variants of PDZ3 contained 3-5 substitutions for 1R, Deuterostomia, Protostomia and Bilateria PDZ3, at positions outside the canonical binding groove (Fig. 1a). The primary structure of AltAll for the ancestral Hexapoda PDZ3 contained 16 substitutions in comparison to the ML sequence. This is because PDZ3 at the Hexapoda node is resurrected from relatively few sequences and from species with a large number of residue substitutions. However, all substitutions were situated outside of the canonical binding groove.

Overall, the AltAll and ML PDZ3 domains showed a similar binding affinity for Eukaryota and native CRIPT (Supplementary Fig. 6 and Supplementary Table 1). Furthermore, secondary structure monitored by circular dichroism were similar for ML and AltAll variants, showing that the folding is robust (Supplementary Fig. 7a-e). We determined the global stabilities by urea denaturation for all AltAll variants (Supplementary Fig. 7f-j, and Supplementary Table 2) and found that they were similar to the corresponding ML variants for 1R, bilaterians and protostomes. The AltAll PDZ3 variant for the ancestor of deuterostomes displayed slightly lower stability compared to the ML variant (Supplementary Fig. 7j and Supplementary Table 2). The curve fitting of AltAll and ML PDZ3 from the ancestor of hexapods suffered from sloping baselines and a short denatured baseline resulting in poorly defined *m*_D-N_ values (Supplementary Fig. 7h). However, qualitatively, it is clear that the AltAll variant is less stable than ML, but well folded. Thus, all AltAll variants were properly folded as judged by the CD spectra (Supplementary Fig. 7a-e) and shape of the urea denaturation curves (as reflected in the *m*_D-N_ values) (Supplementary Fig. 7f-j and Supplementary Table 2).

In conclusion, the AltAll and ML proteins displayed similar properties with regard to stability and binding, and therefore other proteins in the ensemble of likely variants will most probably do that too, including the true ancestral PDZ3 variants (Eick et al. 2017).

### Neuroligin with type 1 PBM is common among deuterostome but not protostome animals

The low affinity of the PDZ3:CRIPT interaction in nematodes and insects inspired us to look at another proposed biological ligand of DLG4 PDZ3, namely neuroligin. Similarly to CRIPT, neuroligin interacts with PDZ3 via its C-terminus at synaptic junctions (Jeong et al. 2019; Irie et al. 1997). To compare the PBMs of neuroligin to those of CRIPT, the NCBI protein sequence database was searched for homologs of *H. sapiens* (UniProt Q8N2Q7) and *C. elegans* (UniProt Q9XTG1) neuroligin-1, as described for CRIPT in the Methods section. *H. sapiens* and *C. elegans* neuroligin sequences were used for the search, because they were both of high quality and represented neuroligin sequences from deuterostomes and protostomes, respectively. The search resulted in 1725 sequences in 437 species – 419 metazoan species and 18 fungal species. Among the neuroligin sequences, type 1 PBM, like the one of CRIPT, was found to be predominant in deuterostomes, while type 2 PBM (ϕ-X-ϕ-COOH) was predominant in protostomes, especially hexapods. Thus, we observe an even more dynamic evolution of the PBM in neuroligin as compared to CRIPT. It appears likely that bilaterian neuroligins recognize different PDZ domains in different species, while most of the neuroligin sequences from fungi and cnidarians – the only species outside of bilaterians that have neuroligin homologs – lack a PBM. However, the lower number of high quality neuroligin sequences makes these results less conclusive than for CRIPT.

## Discussion

We have used ancestral sequence reconstruction and resurrection to better understand the connection between structure, function and evolution of the PDZ3:CRIPT interaction and its biological function. Already at the time of the last common ancestor of all extant metazoa, CRIPT contained a type I PBM with high affinity to most present day DLG PDZ3 domains, and likely to many other PDZ domains with type I PBM specificity. This affinity has remained constant for eons among diverging lineages and until today. But, at the beginning of the evolutionary lineage leading to present day arthropods (approximately 550-600 Myr) a point mutation in the C-terminal residue of CRIPT resulted in substantially decreased affinity for DLG PDZ3, and again, likely all PDZ domains with type I specificity. What is the role of CRIPT and how would this mutation influence function? Large regions of CRIPT are highly conserved among extant species (Supplementary Figure 9) and mutation in CRIPT is rare among humans, and associated with rare Mendelian disorders (Leduc et al. 2016). A recent study including *in vivo* experiments indicated an essential role for CRIPT in dendritic growth (Zhang et al. 2017). Experiments were performed in the nematode *C. elegans*, which, like *D. melanogaster*, has a Thr at the C-terminus of CRIPT. A severely mutated CRIPT (deletion of around 130 residues) resulted in abnormalities in dendritic growth, which could be rescued by expression of *H. sapiens* CRIPT. It was not possible to knock down CRIPT in mouse suggesting a non-redundant and critical cell biological function in vertebrate species (Zhang et al. 2017). On the other hand, it has been possible to knock down one or several of the DLG family members because these proteins can compensate for each other (Bonnet et al. 2013; Howard et al. 2010). In light of their experiments, and the lack of a type I PBM in the C-terminus of CRIPT from *C. elegans* and *D. melanogaster*, Zhang et al suggested that the critical biological action of CRIPT could be partially independent of the PDZ3:CRIPT interaction in these animals. In the present paper we corroborate this idea by showing that PDZ3 domains from extant species of nematode and insect phyla bind their native CRIPT with significantly lower affinities than ancestral PDZ3:CRIPT pairs and those from extant vertebrates, even when CRIPT contains a large hydrophobic residue at the C-terminus, such as Leu in *L. loa* CRIPT. (There are also no obvious non-canonical internal PBMs in CRIPT.) The fact that PDZ3 domains from nematodes and insects have retained the affinity for type I PBM reinforces this notion.

By necessity, many protein interaction studies are performed with domains cut out from highly modular, often large proteins, which are hard to work with in pure form. However, effects from regions outside the binding surface or domain are increasingly considered in protein-protein interaction studies. Indeed, extensions of PDZ domains are associated with several potential functions: 1) protein dynamics-based modulation of target binding affinity, 2) provision of binding sites for macro-molecular assembly, 3) structural integration of multi-domain modules, and 4) expansion of the ligand-binding pocket (Wang et al. 2010). We analyzed the α3 extension in PDZ3 that possibly extends the ligand binding surface beyond the canonical binding groove. Interestingly, while this α3 extension had only minor effects on the low affinity native PDZ3:CRIPT interactions (*L. loa*, *D. melanogaster a*nd *D. ponderosae*), it indeed had a significant effect on binding to the high affinity ancestral eukaryote CRIPT peptide to *L. loa* PDZ3, demonstrating high sequence dependence in the interaction, upon addition of α3. The sequence dependence was also reflected in the stabilities of PDZ3 with and without α_3_, where only *L. loa* PDZ3 displayed a clear stabilization in presence of the helix. With this in mind we note that PDZ3 stabilities vary across present day animals and that ancestral versions of PDZ3 were likely a little more stable. However, all PDZ3 variants were well folded and thus fit for its purpose, binding to another protein. Slow destabilization of domains over evolutionary time is consistent with random mutations, neutral for function and not detrimental as long as folding is correct and stability sufficient.

## Materials and Methods

To collect protein sequences, the NCBI database (NCBI Resource Coordinators) was searched for CRIPT and DLG sequences using search terms and the BLAST tool (Altschul et al. 1990) with the *H. sapiens* DLG4 PDZ3 sequence. Sequences were then aligned using Clustal Omega software (Sievers et al. 2011) and filtered. Non-redundant sequences with information on species of origin, homologous to *H. sapiens* DLG4 PDZ3 or *H. sapiens* CRIPT, respectively, were kept. The final sets of sequences were aligned using Clustal Omega and manually curated. The resulting alignments were used to infer the ancestral sequences of DLG family PDZ3 domains and of CRIPT. For this sequence reconstruction, species trees were downloaded using CommonTree tool from Taxonomy Browser in the NCBI database. For the sequences, which are identical for several species, the lowest common taxonomic rank was used in the tree. The taxonomy trees were modified before the ancestral reconstruction: For every clade branching out into more than 2 daughter clades on one branch, clades were split into smaller clades that only had two daughter clades at the time due to software requirements. The PDZ3 tree was adjusted according to DLG family homology; due to evidence of genome duplication in early chordates, vertebrate sequences were split into DLG1, DLG2, DLG3 and DLG4 clades (Putnam et al. 2008) (McLysaght, Hokamp, and Wolfe 2002). In the resulting CRIPT and PDZ3 trees, species names were replaced with sequence names to match the identifiers in the alignment. The RAxML software (Stamatakis 2014) was used to infer ancestral sequences for PDZ3 and CRIPT. Because of low posterior probabilities for some positions in the reconstructed ML sequences, we reconstructed AltAll sequences for each ML sequence. In the AltAll sequences the amino acid residues with the second highest probabilities were included for each position where the probability of the ML residue was below 0.8.

### Protein expression and Purification

cDNA encoding DLG PDZ3 from the following extant species were cloned into a modified pRSET vector (Invitrogen) and transformed into *Escherichia coli* BL21(DE3) pLys cells (Invitrogen) for expression: *H. sapiens* (DLG4), *S. kowalevskii*, *D. melanogaster*, *D. ponderosae*, *L. loa*, *O. bimaculoides* and *H. vulgaris*. Similarly, reconstructed cDNA encoding the following ancestral DLG PDZ3 (both ML and AltAll variants) were expressed: 1R, Deuterostomia, Bilateria, Hexapoda and Protostomia. The expression construct contained an N-terminal 6xHis-tag. Cells were grown in LB medium at 37°C and overexpression of protein was induced with 1 mM Isopropyl-β-D-thiogalactopyranoside overnight at 18°C. Cells were harvested by centrifugation at 4°C and the pellet re-suspended in 50 mM Tris pH 7.8, 10 % glycerol and stored at −20°C. Pellets were thawed and sonicated 2 × 4 min followed by centrifugation. The supernatant was filtered and added to a pre-equilibrated (50 mM Tris, pH 7.8, 10% glycerol) Nickel Sepharose Fast Flow column (GE Healthcare). Proteins were eluted with 250 mM imidazole and dialysed into 50 mM Tris pH 7.8, 10% glycerol buffer overnight. Proteins were then loaded onto a Q sepharose column for further purification and eluted with a 500 mM NaCl gradient in 50 mM Tris, pH 7.8, 10% glycerol. Protein purity and identity were analyzed by SDS-PAGE and MALDI-TOF mass spectrometry.

### Circular dichroism and urea denaturation experiments

Experiments were performed in 50 mM sodium phosphate, pH 7.45, 21 mM (*I* = 150) at 10°C on a JASCO J-1500 spectrapolarimeter. Far-UV spectra for all PDZ3 domains were recorded from 260 nm to 200 nm with 10 μM protein (the average of 16 scans). Thermodynamic stability of PDZ3 domains was determined by urea denaturation (0 −8.1 M with 0.3 M steps) and monitoried by molar ellipticity at 222 nm with 22 μM PDZ3. Data were fitted to a two-state unfolding process using both free fitting of each parameter or a shared *m*_D-N_ to obtain the midpoint (denoted [Urea]_50%_, where 50% of the protein is folded).

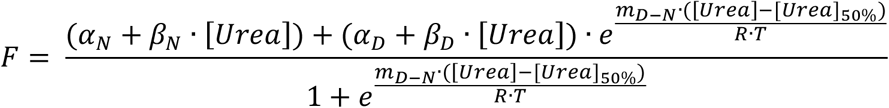

F is the observed spectroscopic signal; α_N_ is the signal of the native state in 0 M urea; β_N_ is the slope of the native baseline; α_D_ and β_D_ are the corresponding parameters for the denaturated state (Clarke et al. (1993)).

### Kinetic experiments

All kinetic experiments were performed in 50 mM sodium phosphate, pH 7.45, 21 mM KCl (I = 150) at 10°C. Binding and dissociation experiments were performed at SX-17 MV stopped-flow spectrometer (Applied Photophysics) and carried out as described previously (Chi et al. (2010)). Briefly, the association rate constant (*k*_on_) was obtained from binding experiments. PDZ3 (1 μM) was mixed rapidly with dansyl labelled (D) CRIPT peptide (concentration range: 2 to 20 μM). Excitation was done at 345 nm and emission was recorded above 420 nm using a long-pass filter). The change in fluorescence signal obtained from D-CRIPT was monitored over time and fitted to a single exponential function to deduce the observed rate constant (*k*_obs_). Observed rate constants were plotted against the corresponding D-CRIPT concentration. Data were fitted to an equation for a reversible bimolecular association reaction (Malatesta 2005) to extract the association rate constant without considering pseudo first order conditions.

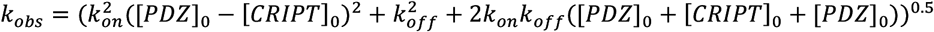

The dissociation rate constant (*k*_off_) was obtained from a separate displacement experiment. An equilibrium complex was pre-formed between PDZ3 and D-CRIPT (2:10 μM) and mixed rapidly with high excess of unlabeled CRIPT (100, 150 and 200μM, respectively). The change in fluorescence signal obtained from dansyl-labelled CRIPT was monitored over time and fitted to a single exponential function to deduce the observed rate constant (*k*_obs_). The average of the three *k*_obs_ values is reported as the dissociation rate constant *k*_off_.

### ITC Binding experiments

All isothermal titration calorimetry (ITC) binding experiments were performed in 50 mM sodium phosphate pH 7.45, 21 mM KCl (I = 150) at 25°C on an iTC200 microcalorimeter (Malvern Instruments). PDZ3 and CRIPT peptide were dialysed against the same buffer to minimize artifacts from buffer mismatch in the titrations. CRIPT peptide was titrated 16 times into the cell containing PDZ3. For every titration the heat release decreased from the binding of CRIPT to PDZ3, due to complex formation and PDZ3 saturation. Change in heat over time was integrated to kcal/mol and fitted to a one site binding model to obtain binding stoichiometry (n), equilibrium binding constant (K_d_) and the enthalphy change upon binding (ΔH).

## Acknowledgements

We thank Eva Andersson for technical assistance. This project was funded by the European Union’s Horizon 2020 research and innovation programme under the Marie Skłodowska-Curie grant agreement No 675341 (to T.G. and P.J.) and by the Swedish Research Council grant no. 2016-04965 (to P.J.).

## Author contributions

L.L. and P.J. conceived and designed the project. L.L performed all experiments and analyzed the data. J.Č. performed all the bioinformatic work including the ancestral sequence reconstruction. L.L., J.Č., and P.J. wrote the initial draft. All authors contributed to preparation of the paper and interpretation of data.

## Data availability

The authors declare that data supporting the findings of this study are available within the paper and its supplementary information file, including probability tables for reconstruction of CRIPT and PDZ3 sequences. Additional raw data are available from corresponding authors upon request.

