## Supplementary material for "Divergent evolution of a protein-protein interaction revealed through ancestral sequence reconstruction and resurrection"

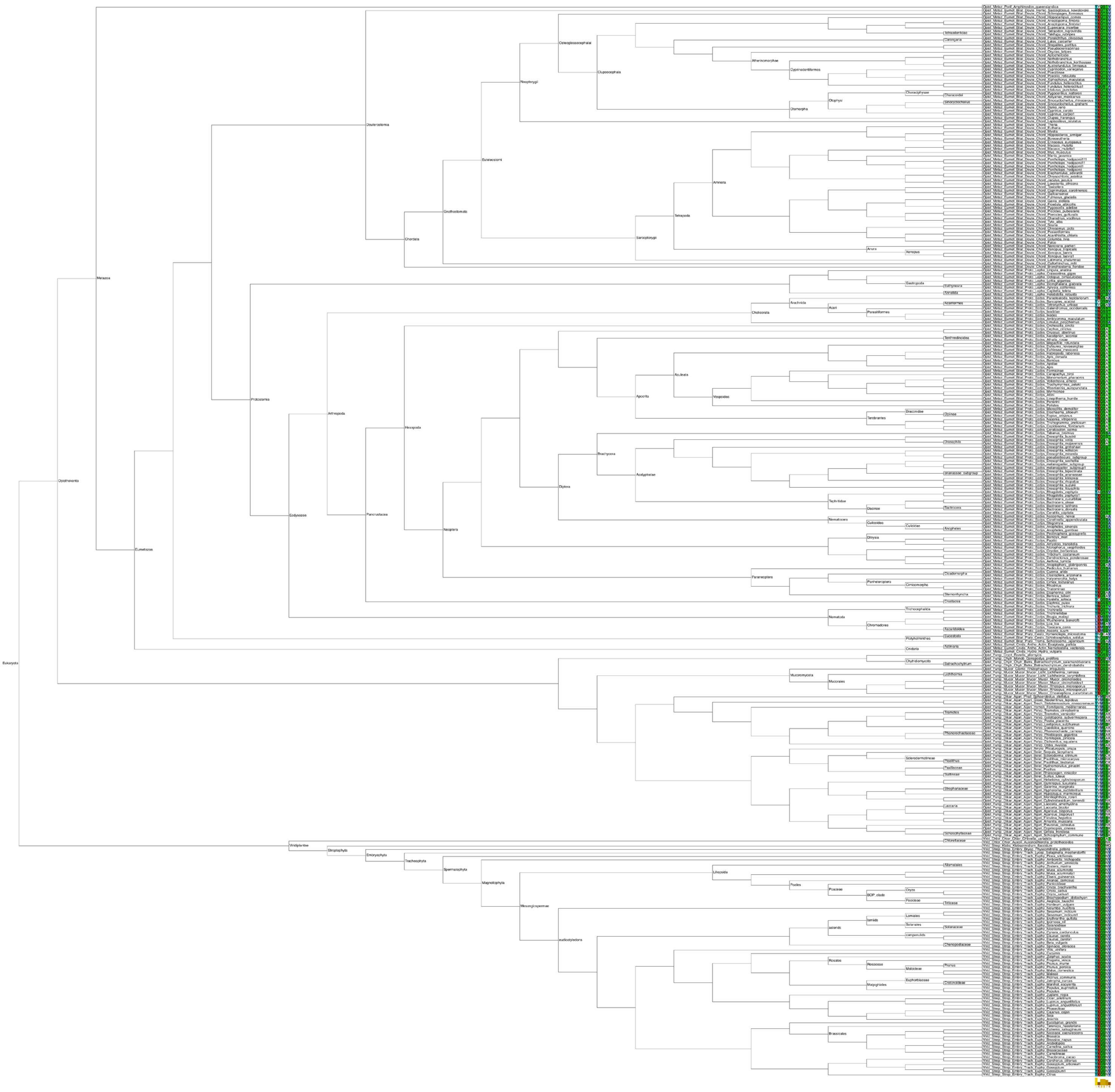

Supplementary Fig. 1
Sequence alignment of all CRIPT sequences used for ancestral reconstruction. The details on sequence conservation are displayed. Conserved amino acids are colored according to their physicochemical properties. The alignment was visualised using Jalview and Dendroscope software (Huson and Scornavacca 2012).

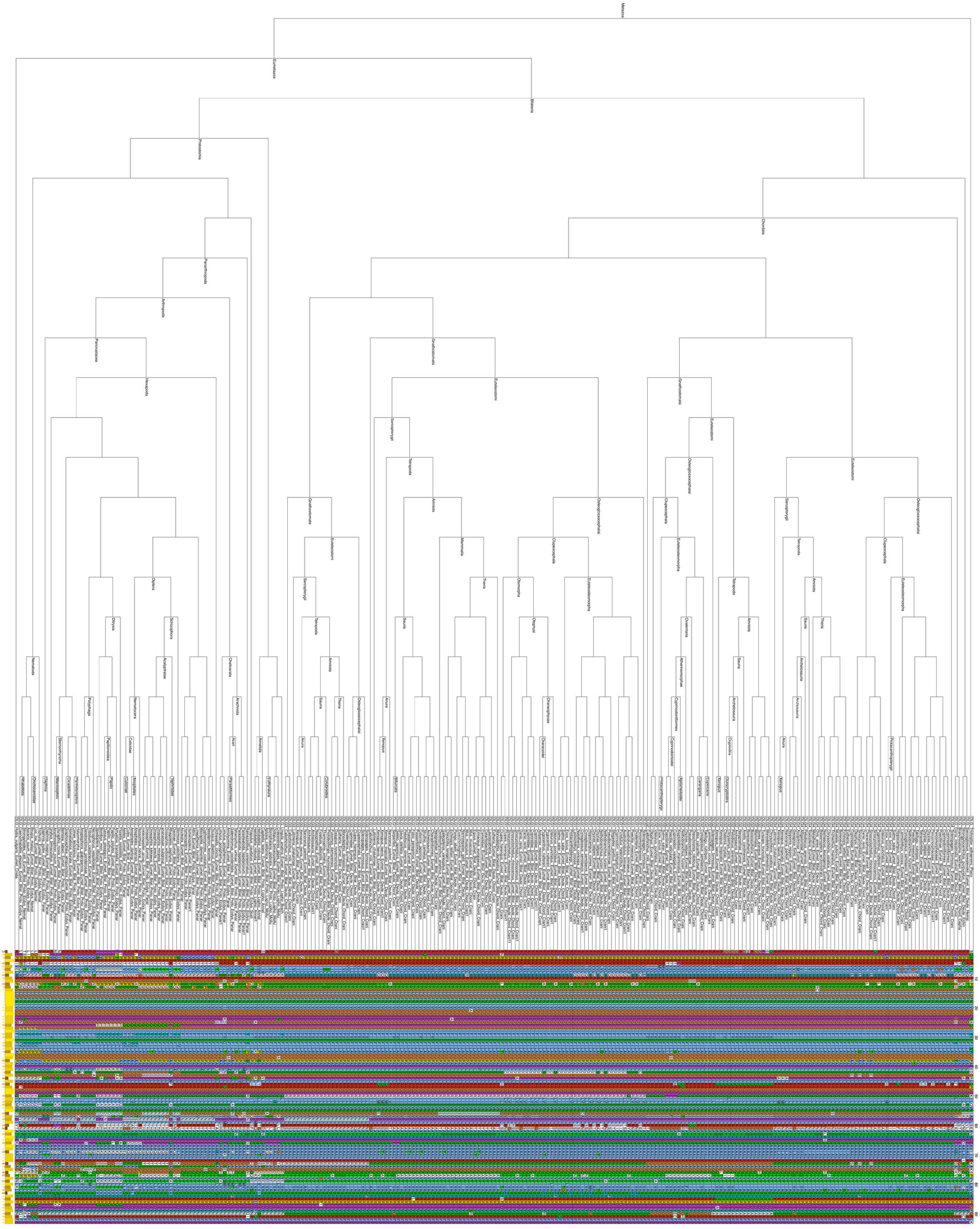

Supplementary Fig.2
Sequence alignment of all DLG PDZ3 sequences used for ancestral reconstruction. Conserved amino acids are colored according to their physicochemical properties. The alignment was visualised using Jalview and Dendroscope software (Huson and Scornavacca 2012).

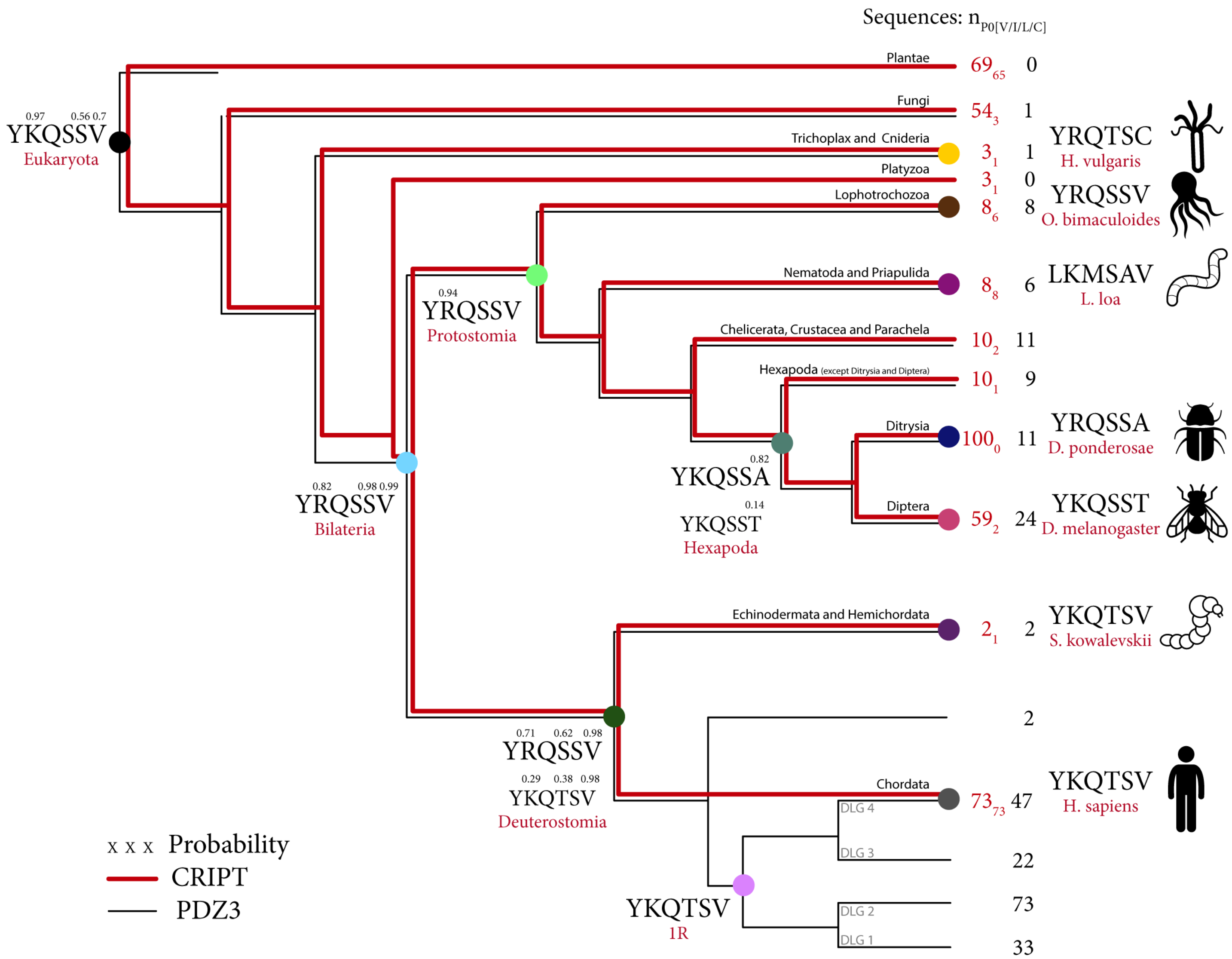


Supplementary Fig. 3
Evolution of the PDZ-binding motif in CRIPT. Simplified species tree with the selected extant and reconstructed ancestral CRIPT C-terminal sequences included in the study. The color code used for the respective evolutionary node is used throughout the manuscript. The posterior probability for each reconstructed CRIPT residue is reported above the residue if different from 1. The red numbers at the branch tips correspond to the number of extant CRIPT sequences included in the ancestral reconstruction for the different animal groups. The red subscript is the number of CRIPT sequences with Val, Ile, Leu or Cys at $P_0$ (thereby possibility for a type I motif binding). The black numbers at branch tips correspond to the number of PDZ3 DLG sequences included in the ancestral reconstruction for the different animal groups.

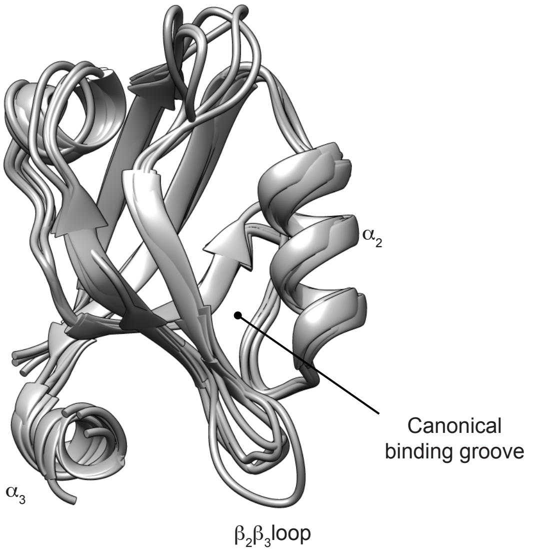


Supplementary Fig. 4
Ribbon diagrams of homology models of extant and ancient variants of PDZ3: Superimposition of homology models for all ancient and extant PDZ3 domains included in the study. The models were predicted from available structures by template search in the PDB database using the SWISS-MODEL software (Waterhouse et al. 2018). Listed are the PDB files used as template for the SWISS-MODEL modelling of the respective PDZ3 domain. *H. sapiens* (PDB:5jxb), *D. melanogaster* (PDB:1um7), *D. ponderosae* (PDB:1um7), *L. loa* (PDB:1pdr), *S. kowalevskii* (PDB:1pdr), *H. vulgaris* (PDB:5jxb), *O. bimaculoides* (PDB:1pdr), Protostomia (PDB:1um7), Bilateria (PDB:1um7), Hexapoda (PDB:1um7), Deuterostomia (PDB:5hed), 1R (PDB:1um7).

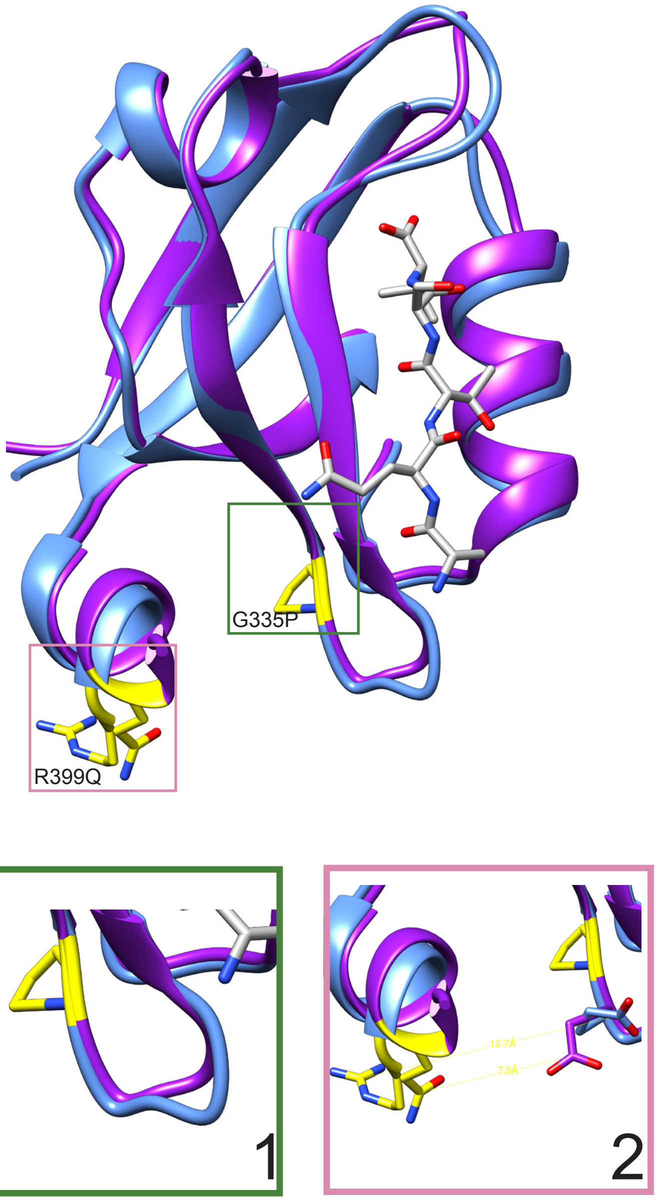


Supplementary Fig. 5
Homology model of *L. loa* PDZ3. Structural comparison of predicted SWISS model of PDZ3 from *L. loa* (purple) and Bilateria (light blue) gives a structural explanation of the relatively weak affinity of *L. loa* PDZ3 for the Eukaryota CRIPT peptide. 1) *L. loa* PDZ3 has a proline instead of a glycine residue in the $\beta_2\beta_3$ loop close to the binding pocket, residues are shown (yellow) 2) The predicted saltbridge between R399 and E334 in the bilaterian complex (distance 12.7 Å) is not present in *L. loa* due to Q399 (yellow). *L. loa* PDZ3 P335G and Q399R were experimentally tested.

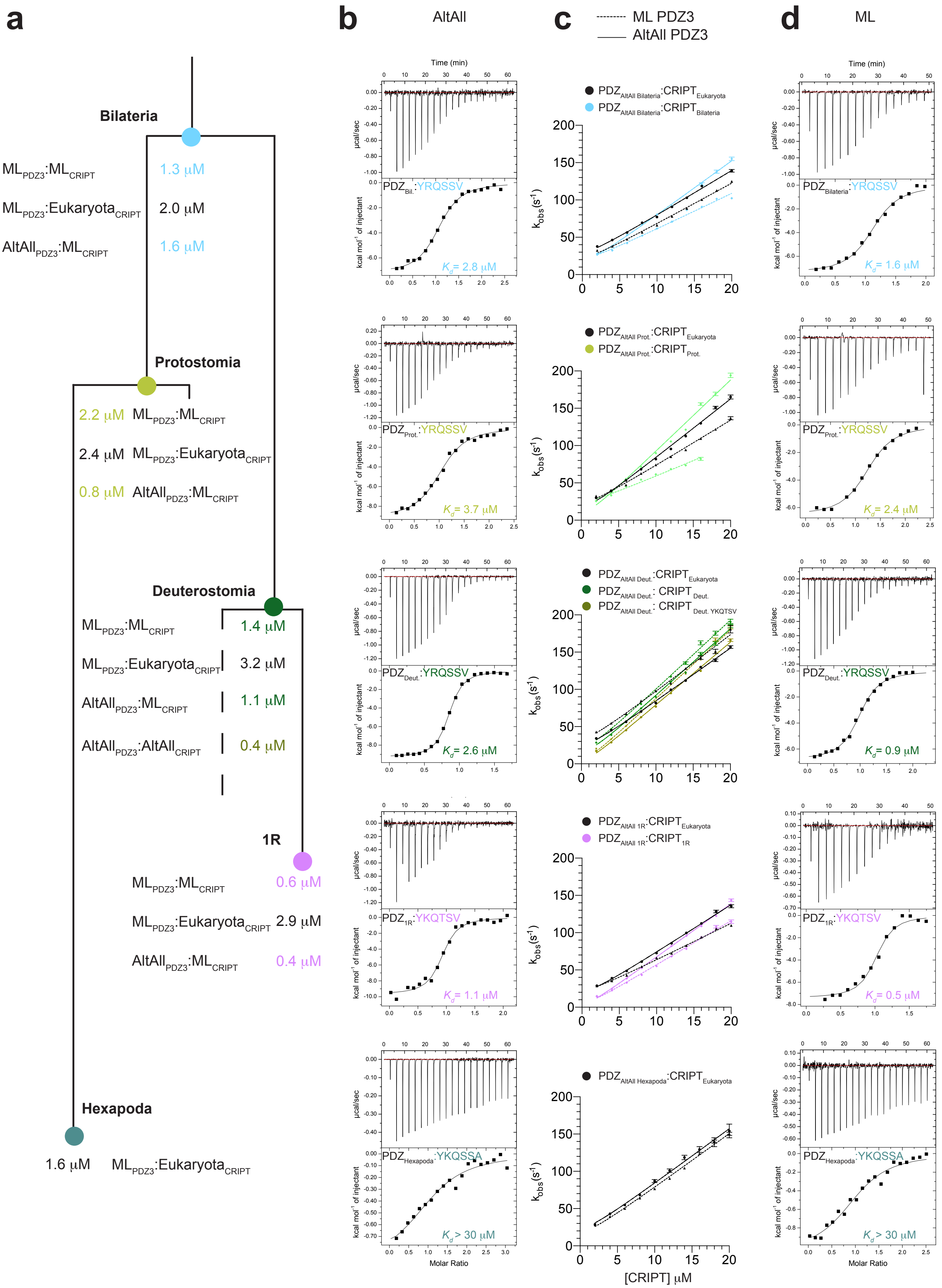

a
b
AltAll
c
ML PDZ3
AltAll PDZ3
d
ML
Bilateria
1.3 μM
2.0 μM
1.6 μM
Protostomia
2.2 μM
2.4 μM
0.8 μM
Deuterostomia
1.4 μM
3.2 μM
1.1 μM
0.4 μM
1R
0.6 μM
2.9 μM
0.4 μM
Hexapoda
1.6 μM
Time (min)
μcal/sec
kcal mol-1 of injectant
Molar Ratio
PDZBil.:YRQSSV
Kd= 2.8 μM
PDZProt.:YRQSSV
Kd= 3.7 μM
PDZDeut.:YRQSSV
Kd= 2.6 μM
PDZ1R:YKQTSV
Kd= 1.1 μM
PDZHexapoda:YKQSSA
Kd > 30 μM
PDZBilateria:YRQSSV
Kd= 1.6 μM
Kd= 2.4 μM
Kd= 0.9 μM
Kd= 0.5 μM
kobs(s-1)
[CRIPT] μM

Supplementary Fig. 6
Comparison of PDZ3:CRIPT interaction from ML and AltAll variants suggest robustness of conclusions to uncertainty in primary structure. Validation of the affinity of ancestral PDZ3:CRIPT interactions by comparison of reconstructed ML and AltAll variants. **a** Schematic diagram of species tree showing the evolutionary nodes of resurrected ancestral PDZ3 and CRIPT variants. The affinity determined by stopped-flow experiments are listed for the following complexes: ML PDZ3:ML CRIPT, ML PDZ3:Eukaryota CRIPT, AltAll PDZ3:ML CRIPT and AltAll PDZ3:AltAll CRIPT and colour coded according to type of CRIPT: reconstructed ancestral variant (ancestor colour code) or Eukaryota (black). **b** Isothermal titration calorimetry experiments for the AltAll PDZ3:CRIPT interactions at the respective nodes. **c** Binding kinetics for ancestral ML and AltAll PDZ3:CRIPT interactions. Observed rate constant were plotted as a function of CRIPT concentration at a constant concentration of PDZ3 (1μM). The colour code represents the type of CRIPT: ML, AltAll, or Eukaryota CRIPT (black), whereas line style represents ML PDZ3 (dashed line) or AltAll PDZ3 (solid line). **d** Isothermal titration calorimetry experiments for the PDZ3:CRIPT interaction in the respective evolutionary nodes of ancestral variants of ML PDZ3 and native CRIPT. Experiments were performed at 25°C for ITC and a 10°C for stopped-flow.

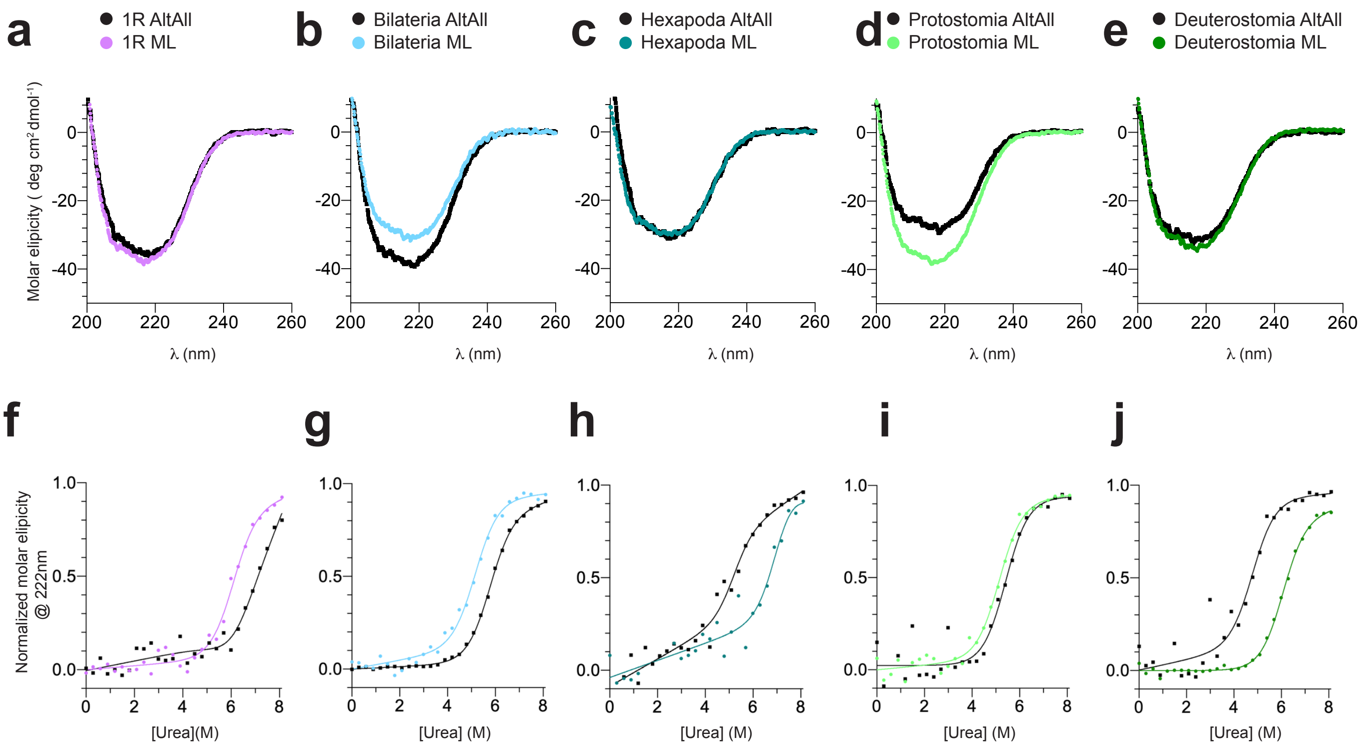


Supplementary Fig. 7
Comparison of structure and global stability of selected ML and AltAll variants of ancestral species. **a-e** Secondary structure content analysed by circular dichroism between 200-260 nm. Each spectrum is an average of 5 individual scans measured at 10°C in 50 mM sodium phosphate, pH 7.45, 21 mM KCl (**I** = 150). **f-j** Urea denaturation (0-8.1 M) of ancestral ML PDZ3 variants (colour code) and the corresponding AltAll (black) ancestral PDZ3 variants, as monitored by circular dichroism at 222 nm. Data were fitted to a two-state model for protein (un)folding. See Table 3 for fitted parameters.

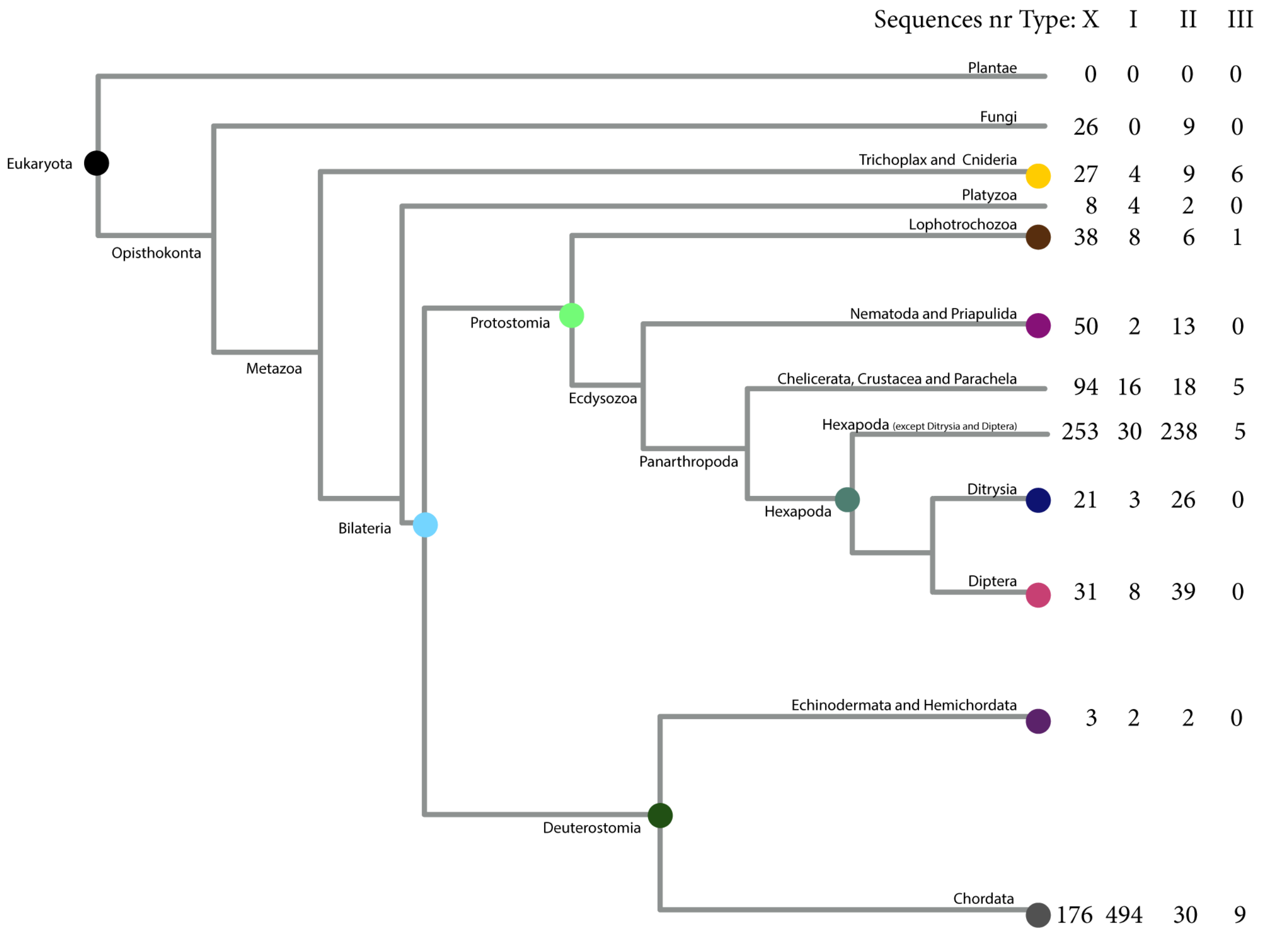


Supplementary Fig. 8
Simplified species tree depicting the evolution of the PDZ-binding motif in Neuroligin. The numbers at the branch tips correspond to the number of extant Neuroligin sequences with type X (not classified), I, II or III motif for the different animal groups.

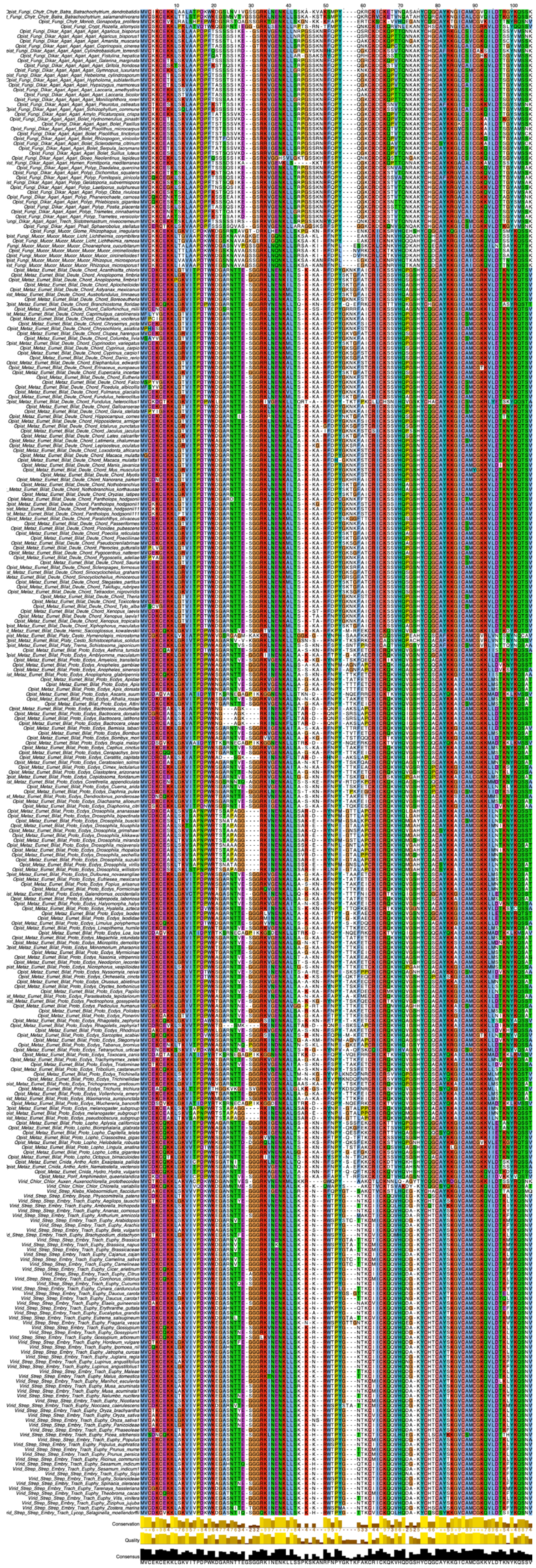

Supplementary Fig. 9
Sequence alignment of all CRIPT sequences used for ancestral reconstruction. The details on sequence conservation and quality are displayed. Conserved amino acids are colored according to their physicochemical properties. The alignment was visualised using Jalview software.

Supplementary Table 2

Comparison of binding constants for the PDZ3:CRIPT interaction from ML and AltAll variants measured by ITC. To facilitate comparison binding parameters for the respective variants of ancestral AltAll (regular) and ML (bold) PDZ3 are listed for different peptide ligands: CRIPT Eukaryota (YKQSSV), CRIPT ML native (YKQTSV, YRQSSV) and AltAll CRIPT for the deuterostome ancestor (the only AltAll CRIPT variant, which is marked with *). Thermodynamic parameters were determined by ITC at 25°C in 50 mM sodium phosphate, pH 7.45, 21 mM KCl (I = 150).

| CRIPT | PDZ3 | $k_{on}$ ($\mu M^{-1}$ $s^{-1}$) | $k_{off}$ ( $s^{-1}$) | $K_d$ ($\mu M$) | |
|---|---|---|---|---|---|
| YKQSSV | 1R | 6.5± 0.1 | 14.1 ± 0.02 | 2.2 ± 0.04 | |
| | **1R ML** | **5.3 ± 0.1** | **15.3 ± 0.3** | **2.9 ± 0.12** | |
| | Deuterostomia | 7.2 ± 0.1 | 15.7 ± 0.1 | 2.2 ± 0.04 | |
| | **Deuterostomia ML** | **7.4 ± 0.2** | **23.6 ± 0.03** | **3.2 ± 0.07** | |
| | Hexapoda | 7.4 ± 0.2 | 11.6 ± 0.1 | 1.6 ± 0.05 | |
| | **Hexapoda ML** | **7.3 ± 0.1** | **11.5 ± 0.1** | **1.6 ± 0.04** | |
| | Protostomia | 7.9 ± 0.2 | 12.0 ± 0.1 | 1.5 ± 0.04 | |
| | **Protostomia ML** | **6.5 ± 0.1** | **15.5 ± 1.3** | **2.4 ± 0.23** | |
| | Bilateria | 6.2 ± 0.1 | 20.7 ± 0.1 | 3.3 ± 0.09 | |
| | **Bilateria ML** | **5.9 ± 0.1** | **11.9 ± 0.3** | **2.0 ± 0.07** | |
| YKQTSV | 1R | 7.2 ± 0.1 | 2.9 ± 0.02 | 0.4 ± 0.01 | |
| | **1R ML** | **6.2 ± 0.1** | **3.5 ± 0.02** | **0.6 ± 0.01** | |
| | Deuterostomia | 8.4 ± 0.1 | 3.3 ± 0.01 | 0.4 ± 0.01 | * |
| | **Deuterostomia ML** | **9.3 ± 0.2** | **5.0 ± 0.03** | **0.5 ± 0.01** | * |
| YRQSSV | Deuterostomia | 9.0 ± 0.1 | 9.6 ± 0.1 | 1.1 ± 0.03 | |
| | **Deuterostomia ML** | **9.4 ± 0.2** | **13.4 ± 0.1** | **1.4 ± 0.03** | |
| | Protostomia | 9.6 ± 0.3 | 7.6 ± 0.05 | 0.8 ± 0.03 | |
| | **Protostomia ML** | **4.6 ± 0.3** | **10.3 ± 0.1** | **2.2 ± 0.16** | |
| | Bilateria | 7.4 ± 0.2 | 12.0 ± 0.3 | 1.6 ± 0.07 | |
| | **Bilateria ML** | **5.1 ± 0.2** | **6.8 ± 0.1** | **1.3 ± 0.07** | |

Supplementary Table 1

Comparison of binding constants for the PDZ3:CRIPT interaction for ML and AltAll variants. To facilitate comparison, binding parameters for the respective variants of ancestral AltAll (regular) and ML (bold) PDZ3 are listed for different peptide ligands: CRIPT Eukaryota (YKQSSV), CRIPT ML native (YKQTSV, YRQSSV) and AltAll CRIPT for the deuterostome ancestor (the only AltAll CRIPT variant, which is marked with *). The association rate constant ($k_{on}$) was obtained from binding experiment, and the dissociation rate constant ($k_{off}$) from displacement experiment, by stopped-flow experiments at 10°C in 50 mM Sodium Phosphate, pH 7.45, 21 mM KCl (I = 150). The dissociation constant $K_d$ was calculated as $k_{off}/k_{on}$.

| CRIPT | PDZ3 | ΔH($^{kcal}/_{mol}$) | TΔS($^{kcal}/_{mol}$) | $K_d$ (μM) | |
|---|---|---|---|---|---|
| YKQTSV | 1R | -9.7 ± 0.2 | -1.5 | 1.1 ± 0.2 | |
| | **1R ML** | **-7.6 ± 0.2** | **1.0** | **0.5 ± 0.1** | |
| | Deuterostomia | -9.3 ± 0.1 | -0.9 | 0.7 ± 0.04 | * |
| | **Deuterostomia ML** | **-8.8 ± 0.1** | **-0.6** | **0.9 ± 0.1** | * |
| YRQSSV | Deuterostomia | -7.4 ± 0.1 | 0.2 | 2.6 ± 0.2 | |
| | **Deuterostomia ML** | **-6.8 ± 0.1** | **0.9** | **2.1 ± 0.2** | |
| | Protostomia | -9.2 ± 0.2 | -1.8 | 3.7 ± 0.4 | |
| | **Protostomia ML** | **-6.4 ± 0.2** | **1.3** | **2.4 ± 0.3** | |
| | Bilateria | -7.2 ± 0.1 | 0.3 | 2.8 ± 0.3 | |
| | **Bilateria ML** | **-6.8 ± 0.2** | **1.1** | **1.6 ± 0.3** | |
| YRQSSA | Hexapoda | -1.2 ± 0.2 | 5.0 | 31 ± 7.4 | |
| | **Hexapoda ML** | **-2.3 ± 2.0** | **3.5** | **56 ± 25** | |
| YRQSST | Hexapoda | -0.9 ± 0.2 | 5.2 | 31 ± 7.3 | |
| | **Hexapoda ML** | **-1.1 ± 0.1** | **5.3** | **21 ± 3.7** | |
